# Chromosome fusion and programmed DNA elimination shape karyotypes of parasitic nematodes

**DOI:** 10.1101/2023.12.21.572835

**Authors:** James R. Simmons, Brandon Estrem, Maxim V. Zagoskin, Ryan Oldridge, Sobhan Bahrami Zadegan, Jianbin Wang

## Abstract

A growing list of metazoans undergo programmed DNA elimination (PDE), where a significant amount of DNA is selectively lost from the somatic genome during development. In some nematodes, PDE leads to the removal and remodeling of the ends of all germline chromosomes. In several species, PDE also generates internal breaks that lead to sequence loss and an increased number of somatic chromosomes. The biological significance of these karyotype changes associated with PDE and the origin and evolution of nematode PDE remain largely unknown. Here, we assembled the single germline chromosome of the horse parasite *Parascaris univalens* and compared the karyotypes, chromosomal gene organization, and PDE features among ascarid nematodes. We show that PDE in *Parascaris* converts an XX/XY sex-determination system in the germline into an XX/XO system in the somatic cells. Comparisons of *Ascaris, Parascaris,* and *Baylisascaris* ascarid chromosomes suggest that PDE existed in the ancestor of these parasites, and their current distinct germline karyotypes were derived from fusion events of smaller ancestral chromosomes. The DNA breaks involved in PDE resolve these fused germline chromosomes into their pre-fusion karyotypes, leading to alterations in genome architecture and gene expression in the somatic cells. Cytological and genomic analyses further suggest that satellite DNA and the heterochromatic chromosome arms play a dynamic role in the *Parascaris* germline chromosome during meiosis. Overall, our results show that chromosome fusion and PDE have been harnessed in these ascarids to sculpt their karyotypes, altering the genome organization and serving specific functions in the germline and somatic cells.

## Introduction

Faithful maintenance of genomes is essential for the survival and propagation of species. However, some species undergo dramatic genome changes during development through programmed DNA elimination (PDE).^1-4^ PDE (also known as chromatin diminution) was discovered by Theodor Boveri in the 1880s in the horse parasitic nematode *Parascaris spp*.^5^ Since then, many species from diverse phylogenetic groups have been identified to undergo PDE.^1-4,6-8^ The distinct features of PDE from each group and its sporadic presence in metazoa suggest PDE evolved independently multiple times.^1-3^ The broad distribution of PDE further suggests it plays an important role(s) in divergent species.

Studies in nematodes have provided the most comprehensive understanding of PDE in metazoa.^8,9^ Cytological, molecular, and genomic studies in ascarids^10^ (specifically, *Ascaris*^11^ and *Parascaris*) have identified several features of nematode PDE, including 1) the elimination of genes and repeats during early embryogenesis;^12-14^ 2) the consistent breakage of chromosomes at specific loci in all pre-somatic cells;^12-14^ 3) the selective segregation of retained DNA;^15,16^ 4) the healing of DNA breaks with telomere addition;^17,18^ and 5) the removal of all germline chromosomes ends.^14,16^ Recently, several of these features of PDE were also observed in the free-living nematode *Oscheius tipulae*^19,20^ as well as other *Oscheius spp., Caenorhabditis, Auanema*, and some *Mesorhabditis* species.^21^ However, PDE in the nematode *Strongyloides* is divergent,^9,22^ and it occurs during mitotic oocyte division and functions in sex determination.

Significant differences in the mechanisms and consequences of PDE exist between the parasitic ascarids and the free-living *O. tipulae*. The amount of eliminated DNA ranges from 0.6% in *O. tipulae* to 18% in *Ascaris* and 90% in *Parascaris*.^8^ The number and expression profiles of the eliminated genes and the sequence requirement for DNA breaks also differ. These differences suggest some functions and mechanisms of PDE are unique in specific nematode groups. Another key difference is karyotype changes. The number of *O. tipulae* chromosomes (5A and 1X; A = autosome and X = sex chromosome) does not change after PDE.^19,20^ However, *Ascaris* PDE converts the 24 (19A and 5X) germline chromosomes into 36 (27A and 9X) somatic chromosomes;^14^ and the single (1A/X) *Parascaris* germline chromosome is split into 35 (defined by cytology) somatic chromosomes.^16^ Comparative genomics has identified seven ancestral nematode chromosomal elements with a set of conserved genes (Nigon elements).^20^ Analyses of these Nigon elements, other orthologous genes, and PDE in *Ascaris* have revealed that the internal breaks (breaks occurring outside of subtelomeric regions) within the *Ascaris* sex chromosomes coincide with sites of apparent chromosome fusion that formed germline chromosomes,^14,20^ suggesting these internal PDE DNA breaks occur at the ends of ancient sex chromosomes. However, it is unclear if the internal breaks in *Ascaris* autosomes also occur near the sites of chromosome fusions or if they are *de novo* breaks that recently arose.^2,8^

The single, large (2.5 Gb) germline chromosome of *Parascaris* enabled early cytological analyses, making it a historical model for chromosome biology.^23,24^ In addition to providing the first observation of PDE, *Parascaris* studies also contributed to the establishment of Sutton-Boveri chromosome theory of heredity and the understanding of meiosis,^25^ centrosome function in chromosome segregation, and the biology of holocentric chromosomes.^26,27^ Here, we used the single, large germline chromosome of the *Parascaris* to study the chromosomal locations and origin of the DNA breaks, their contribution to karyotype changes, and the evolution of PDE in nematodes. We assembled the *Parascaris* germline genome and revealed the mosaic nature of its autosome and sex chromosome regions. Our analyses of germline chromosome organization in *Parascaris* and comparisons of these features with other ascarids revealed that ascarid germline genomes were formed by extensive chromosome fusions. PDE resolves the fused chromosomes and restores them to their pre-fused karyotypes in all somatic cells. Our genomic and cytological analyses of *Parascaris* centromere reorganization and satellite DNA also suggest their potential functions during meiosis. Together, our comparative genomics and evolutionary studies provide new insights into nematode PDE demonstrating its key contribution to chromosome organization and the biology of nematodes.

## Results

### The single *Parascaris* germline X chromosome encompasses both somatic autosomes and sex chromosomes

There are two known species of the horse parasitic nematode *Parascaris, P. univalens* with one pair of germline chromosomes (2n = 2, Figure 1A) and *P. equorum* with two (2n = 4).^28^ *P. univalens* is the main species currently observed in equines^29^ and used in this study. Here we use *P. univalens* and *Parascaris* interchangeably. We previously assembled a draft genome of *Parascaris* with 50% of the assembled sequences in scaffolds (N50) >= 1.83 Mb, enabling comparative analysis of DNA breaks, eliminated genes, and their expression.^13^ However, the genome assembly has > 1,000 scaffolds and thus it did not allow analysis of the karyotype changes between the germline and somatic chromosomes derived from PDE. Here, we used PacBio long reads and Hi-C to assemble the euchromatic region of the 2.5-Gb germline chromosome into a single scaffold of 240 Mb (Figure 1) that constitutes 98.5% of all assembled sequences. The remainder of the genome (2.2 Gb) is mainly composed of pentanucleotide and decanucleotide repeats in the heterochromatic arms^16^ that are eliminated in PDE and could not be assembled despite having generated > 1 GB of long reads over 100 Kb. The germline genome assembly enables the identification of DNA breaks and new telomere addition sites that generate the somatic genome, thus defining the ends of all somatic chromosomes (Figure 1E and see below). Overall, we have assembled the full euchromatic region of the *Parascaris* germline genome and the complete sequence of the somatic genome.

**Figure 1.**
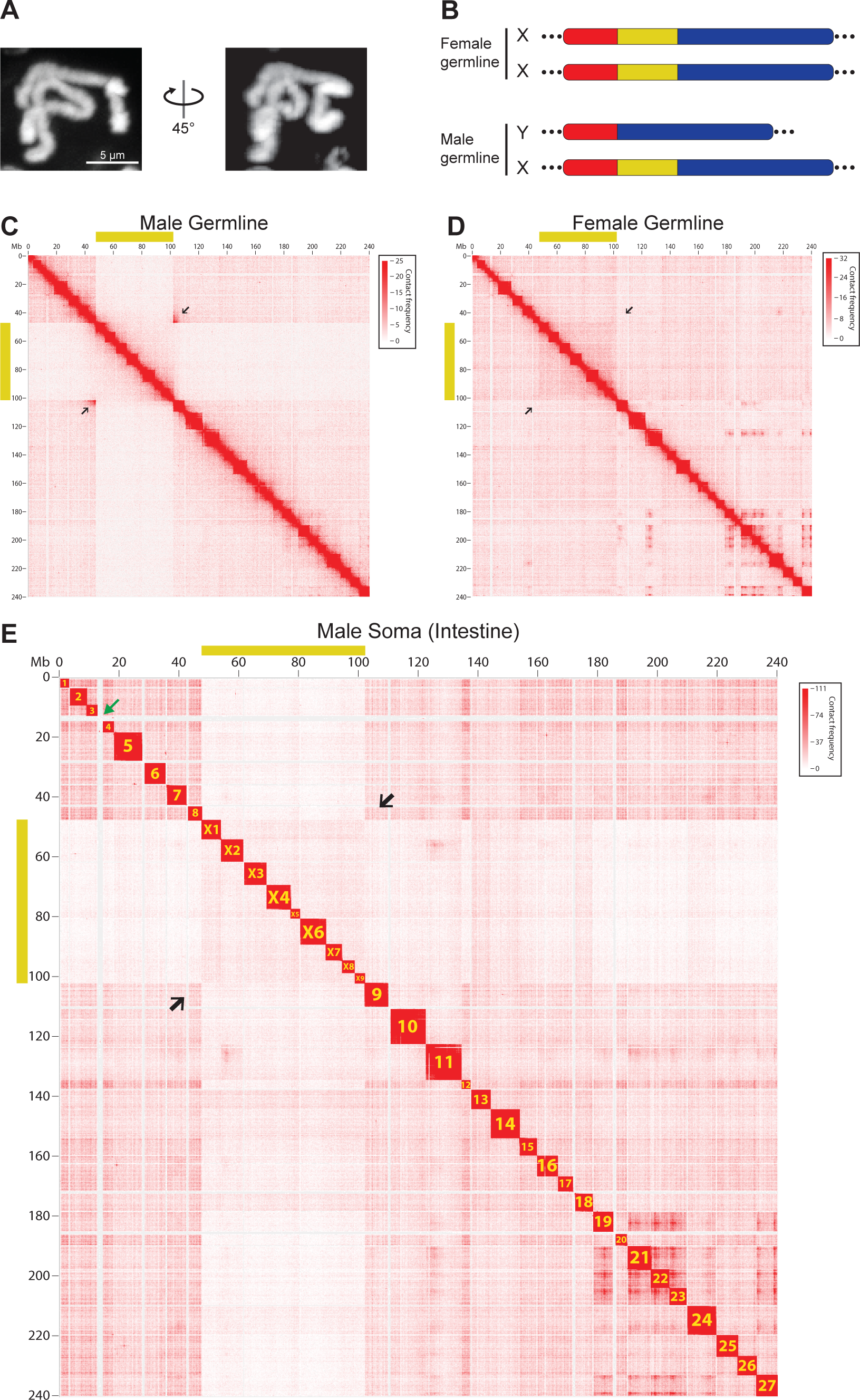
The single *Parascaris* germline X chromosome consists of both somatic autosomes and sex chromosomes. **A.** Hoechst staining of the single, condensed pair of *Parascaris univalens* germline chromosomes from the mitotic region of the testis. **B.** Karyotypes for *Parascaris* germline chromosome derived from this study (see Figure 1C-E). *Parascaris* females have two X chromosomes while males have one X and a shorter Y chromosome that lacks the sex chromosome region (yellow). Autosomal regions are represented in red and blue. Coverage for PacBio long reads spanning the junctions of autosome and sex chromosome regions are shown. Dotted lines at the ends indicate the large heterochromatic arms (90% of the genome) that are mainly comprised of satellite repeats. **C-E.** *Parascaris* Hi-C interaction heatmaps reveal germline karyotypes and their changes after PDE. Yellow bars along the x and y axes indicate the sex chromosome region of the *Parascaris* X chromosome. Black arrows point to the boundaries between autosome and sex chromosome regions. **C.** Reduced Hi-C interactions (∼50%) in the sex chromosome region and strong interactions at the junctions (arrows) suggest two distinct chromosomes (XY) in the male germline (see Figure 1B). **D.** Consistent Hi-C interactions across the regions with no elevated interactions at the junctions (arrows) indicate the same pair of chromosomes (XX) in the female germline. **E.** PDE breaks the germline genome into 36 distinct somatic chromosomes. Shown are Hi-C interactions from the male intestine (see Figure S1 for Hi-C data in the female intestine). The somatic chromosomes were named based on their order in the germline and by their autosome or sex chromosome identities. The green arrow between chromosomes 3 and 4 indicates the largest eliminated region (∼2 Mb) of euchromatic DNA.

Previous cytological work was not able to identify distinct autosome and sex chromosome regions within the single *Parascaris* germline chromosome. Thus, it was also not clear if there were different male and female chromosomes, or how sex is determined in the *Parascaris* chromosomes. Using genome sequencing and Hi-C, we identified a 55 Mb genomic region in the male germline with reduced (50%) overall genome coverage (Figure 2A) and 3D genome interactions (Figure 1C). Comparative sequence analysis indicates this region is orthologous to the *Ascaris* sex chromosomes.^14^ Strong Hi-C interactions between the two ends of this 55 Mb region (arrows in Figure 1C) suggest that these two ends are adjacent to each other at the sequence level in one of the two homologous chromosomes. These strong Hi-C interactions are not seen in the female germline or male somatic cells (arrows in Figure 1D-E), suggesting they are specific to the male germline. These data lead us to propose an XX/XY karyotype (XX for female and XY for male) for the *Parascaris* germline (Figure 1B), with the Y chromosome lacking this 55 Mb sex chromosome region. Somatic genome sequence (Figure 2A) and Hi-C data from males (Figure 1E) and females (Figure S1) further confirm that this 55 Mb region is the origin of somatic sex chromosomes.

**Figure 2.**
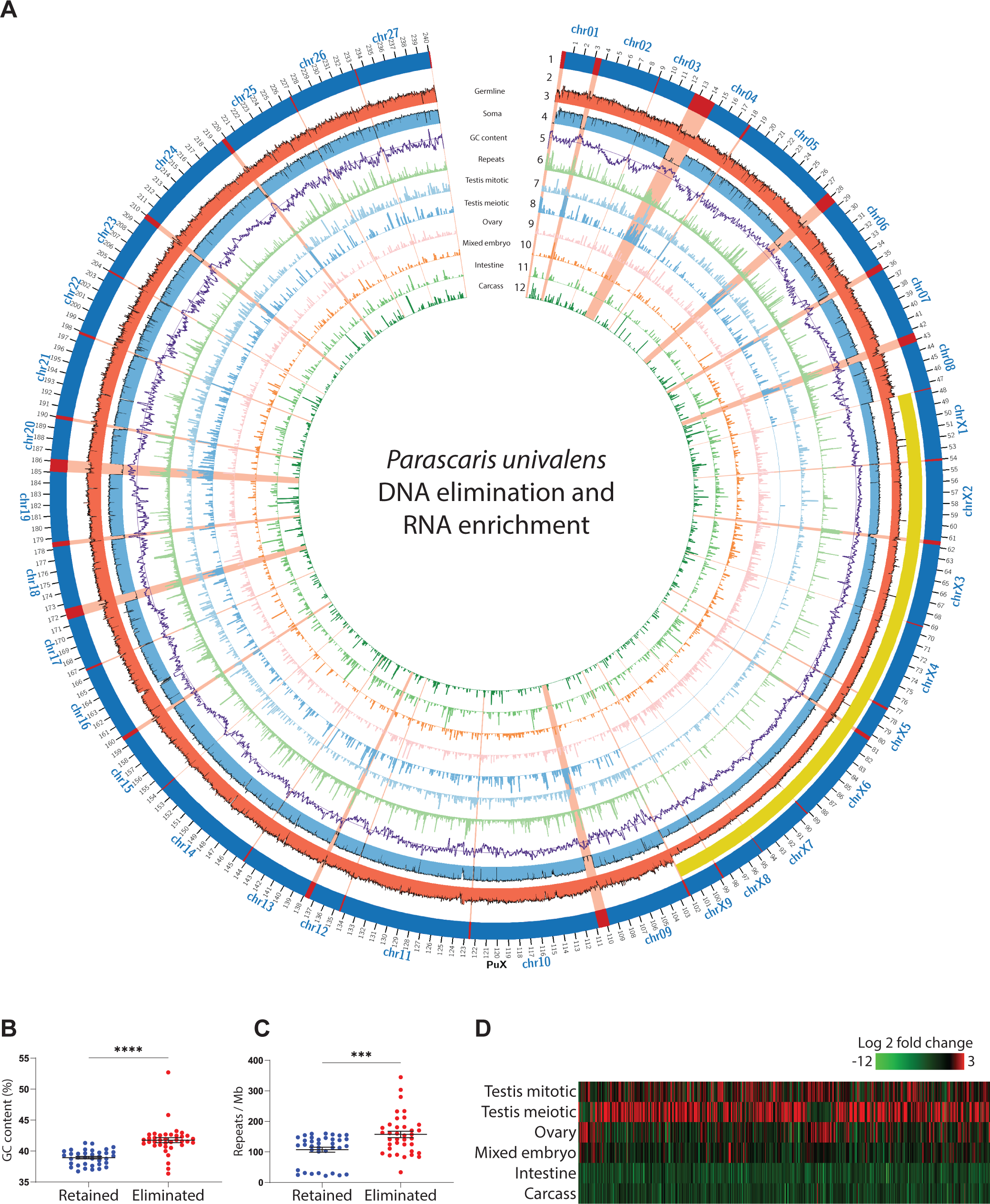
PDE breaks the *Parascaris* germline chromosome and silences germline-expressed genes. **A.** Features of the *Parascaris* X chromosome illustrated in a circos plot. Descriptions of the features from outer to inner circles: 1). Retained (dark blue) and eliminated DNA (red); the eliminated regions are highlighted in red across all circles; 2). The sex chromosome region (yellow bar); 3-4). Genomic (Illumina) read coverage from testis (red) and male intestine (light blue); 5). GC content (purple) and the average GC line; 6). Repeat density (light green) and the average repeat line; and 7-12). Expression (normalized RNA-seq) data for male mitotic region (light blue), male meiotic region (blue), ovary (pink), mixed embryos (orange), intestine (green), and carcass (dark green). **B.** Eliminated regions have higher GC content. **C**. Eliminated DNA contains more repeats. **D**. Eliminated genes are only expressed in the germline - mainly in the testis. The heatmap shows 743 expressed eliminated genes (rpkm >= 5 in at least one of the tissues; see Table S1). P-values for B and C were determined using Welch’s t-tests (*** P < 0.01 and **** P < 0.001).

Through PDE, the single germline X chromosome becomes 36 somatic chromosomes (27A and 9X), as illustrated by 36 distinct Hi-C blocks in the somatic cells (Figures 1E and S1). The number of somatic chromosomes is close to (but not exactly) the reported number (35; 29A and 6X) from an earlier cytological study.^16^ This difference is likely due to the difficulties in visualizing and counting somatic chromosomes using cytology, particularly since seven somatic chromosomes are smaller than 4 Mb. Nevertheless, the junctions of autosome and sex chromosome regions in the germline chromosome correspond to the sites of PDE DNA breaks, allowing a clear resolution and separation of the autosomes and sex chromosomes from the single germline chromosome following PDE. Thus, PDE in *Parascaris* converts the XX/XY sex-determination system in the germline into an XX/XO system (XX for female and XO for male) in the somatic cells. Like *Ascaris*,^14^ no fusion or rearrangement of the chromosome fragments was observed in *Parascaris* following PDE. Notably, while the germline chromosomes of *Ascaris* and *Parascaris* are drastically different, the number of somatic autosomes (27) and sex chromosomes (9) are the same, raising the interesting possibility that the two nematodes may have matching somatic chromosomes (see below).

### PDE breaks the *Parascaris* germline chromosome and silences germline-expressed genes via their elimination

Sequence analysis indicates that ∼12 Mb of DNA is eliminated from the euchromatic genome (Figure 2A). The eliminated DNA is derived from 37 blocks within the germline and the size varies from 10 kb to 2 Mb (average 324 kb). The resulting 36 somatic chromosomes range in size from 2.8 Mb to 11.9 Mb (average 6.3 Mb). We identified 72 regions in the germline genome where DNA breaks and telomere addition occur. These sites are known as chromosomal breakage regions (CBRs), and they are at the junctions of the retained and eliminated regions (Figure 2A). As observed in *Ascaris*, the telomere addition sites within the CBRs are heterogeneous, and no conserved sequence motif or structural features were present within the CBRs, suggesting a potential sequence-independent mechanism for their recognition.^13,14^ The assembled eliminated regions are in general associated with high GC content (Figure 2A-B) and are enriched with non-satellite repeats (Figures 2A and 2C). Overall, the data provides a complete view of the DNA breaks and eliminated sequences along the single germline chromosome.

Using our comprehensive RNA-seq data from 17 developmental stages, we identified 16,603 genes in *Parascaris* (see Methods and Table S1). PDE leads to the elimination of 1,180 (7.1%) genes. Expression analysis revealed these eliminated genes are mostly expressed in the germline, with the majority of them (59.3%) exclusively expressed during spermatogenesis (Figures 2A and 2D, and Table S1). This is consistent with our previous observation in *Ascaris*,^12-14^ suggesting PDE removes and silences testis-specific genes in somatic cells. A comparison between *Ascaris* and *Parascaris* also reveals that orphan genes - genes that lack a detectable homolog in the other species - are more commonly seen in the eliminated regions (47.3%) than in the retained DNA (25.2%). The relaxed constraint for the eliminated DNA supports a hypothesis that eliminated regions can serve as a reservoir where novel genes are better tolerated, and their divergence by rapid mutation promotes the speed of gene selection and evolution.^2,30,31^

### Centromere reorganization facilitates *Parascaris* meiosis and PDE

Early cytological studies revealed that *Parascaris* germline chromosomes are holocentric.^26,27^ The position of kinetochore–microtubules is highly dynamic: during germline mitosis, microtubules attach along the entire chromosome; during meiosis, they are concentrated at the ends of the chromosome; and in early embryos prior to PDE, they are enriched in the euchromatic region of the germline chromosomes.^26,27^ The reorganization of the microtubules to the ends of the chromosomes is believed to circumvent the problem that could arise from random microtubule attachment along the holocentric chromosomes, which may prevent the proper segregation of homologous chromosomes during meiosis.^32-34^ To examine the dynamic centromere and microtubule organization at the molecular level, we used CUT&RUN^35,36^ and immunohistochemistry on the epigenetic mark of centromeres, histone H3 variant CENP-A (also known as CenH3),^37,38^ in germline and early embryos (Figure 3). Our CUT&RUN data showed that CENP-A deposition is inversely correlated with RNA transcription (Figure 3A-B). This is consistent with CENP-A distributions in other nematodes,^15,39^ suggesting the CENP-A CUT&RUN enriched regions represent centromeric regions.

**Figure 3.**
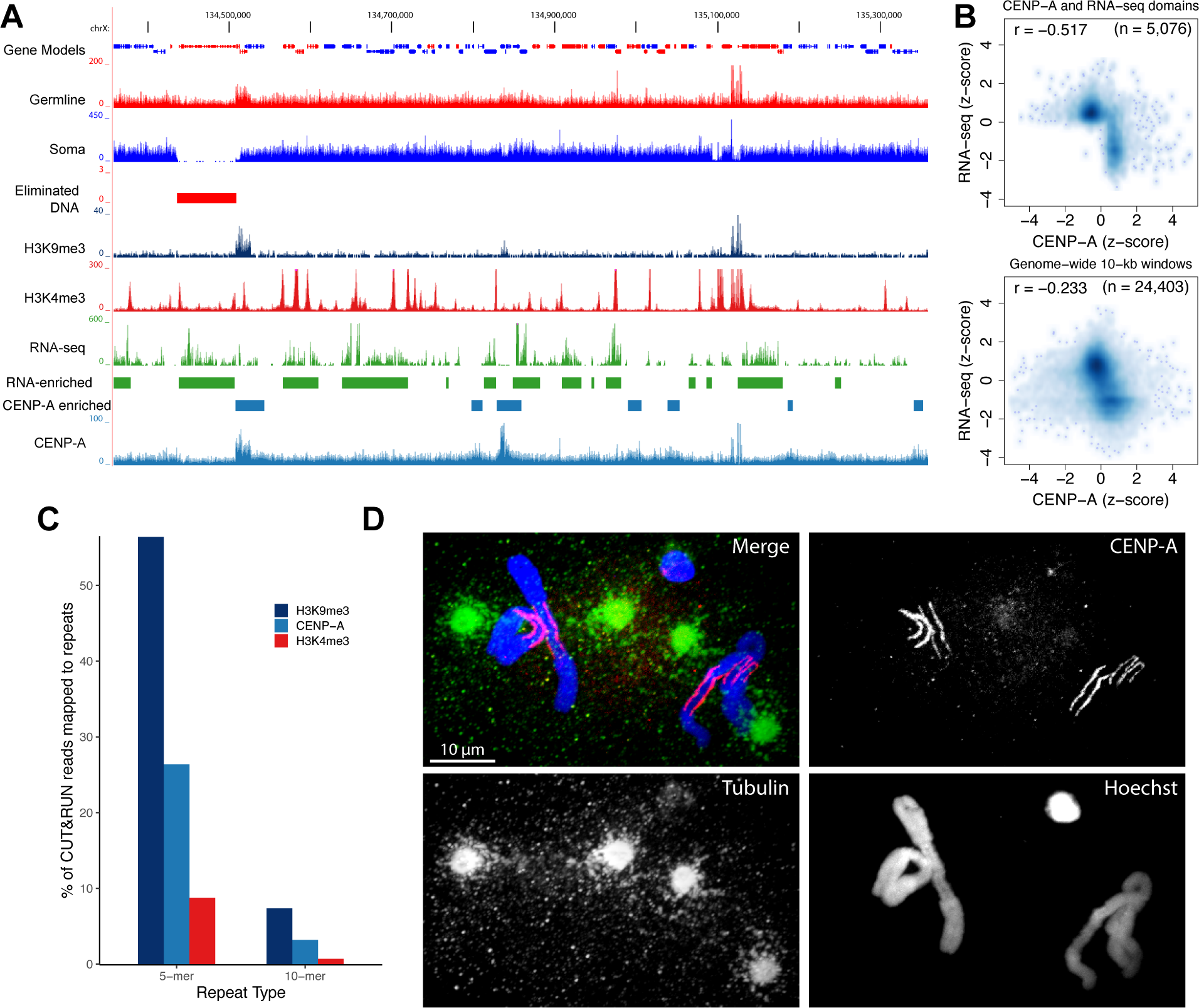
Centromere reorganization facilitates *Parascaris* meiosis and PDE. **A.** CENP-A distribution in the *Parascaris* genome. A browser view of RNA-seq and CUT&RUN data from the ovary tissue for a genomic region of 1 Mb. **B.** Meta-analysis illustrating a genome-wide inverse relationship between RNA transcripts and CENP-A deposition. The top plot is derived from CENP-A and RNA-seq enriched genomic domains (see enriched regions in Figure 3A), showing a strong inverse relationship. The bottom plot is obtained from 10-kb windows for the entire genome showing a modest but consistent negative correlation. **C.** Differential CENP-A, H3K9me3, and H3K4me3 distribution on the pentanucleotide (5-mer) and decanucleotide (10-mer) repeats. **D.** CENP-A staining in early embryos shows CENP-A aligns at the outer side of the chromosomes towards centrosomes. Early embryos were stained with Hoechst (blue) and with antibodies against CENP-A (red) and tubulin (green). Note that CENP-A specifically labels the euchromatic regions of the chromosomes in both the germ cell and the pre-somatic cells. Tubulin staining shows centrosomes on both sides of the dividing chromosomes, with spindle microtubules contacting the CENP-A labeled regions.

The germline heterochromatic chromosome arms are composed of pentanucleotide and decanucleotide repeats. Previous DNA FISH work indicated the ends of the arms were capped by the decanucleotide.^16^ Our CUT&RUN data from the male and female germline (largely consisting of meiotic cells) showed that on average, 26% of CENP-A reads match pentanucleotides, while about 3% of reads correspond to the decanucleotide (Figure 3C), suggesting CENP-A is enriched at the proximal ends of the chromosomes during meiosis. In the 2-cell embryos prior to PDE, immunohistochemistry indicated CENP-A is only at the euchromatic regions in both germline and pre-somatic cells, suggesting CENP-A is no longer in the heterochromatic arms (Figure 3D). This lack of CENP-A at the arms is consistent with the trailing chromosome arms observed during the anaphase in the 2-cell embryos.^26,27^ The lack of CENP-A in the heterochromatic arms also provides a mechanism for their elimination after DNA breakage. Thus, dynamic CENP-A deposition facilitates PDE elimination and may also contribute to the segregation of homologous chromosomes during meiosis by repositioning centromeres to the ends of the holocentric chromosome.^26,27^

### *Ascaris, Parascaris,* and *Baylisascaris* have the same number of syntenic somatic chromosomes

The number of somatic chromosomes is the same between *Ascaris* and *Parascaris* (27A and 9X; see Figures 1E and 2A). Our analysis of available genomic data of another related parasitic nematode of the giant panda, *Baylisascaris schroederi*,^40^ identified eliminated DNA via reduced genome coverage, chromosome end remodeling, and new telomere addition sites, demonstrating that *B. schroederi* also undergoes PDE. Interestingly, *B. schroederi* has the same number (27A and 9X) of somatic chromosomes as *Ascaris* and *Parascaris*. Comparison of these ascarids’ chromosomes enables us to define the synteny among the germline genomes (Figures 4A, S2, and Table S2). Despite a few inversions and rearrangements, most of the three ascarid genomes have perfectly aligned synteny blocks for regions corresponding to the 36 somatic chromosomes (Figure 4B and Table S2). Interestingly, four junctions where somatic chromosomes (chr1/chr2, chrX2/chrX3, chrX6/chrX7, and chr19/chr20) are resolved from within the *Parascaris* germline chromosome show different organization in other ascarid germline chromosomes (Figures 4A and S2). Thus, while the germline karyotypes are drastically different between *Parascaris* (one chromosome) and the other ascarids (24-27 chromosomes), their somatic karyotypes are the same.

**Figure 4.**
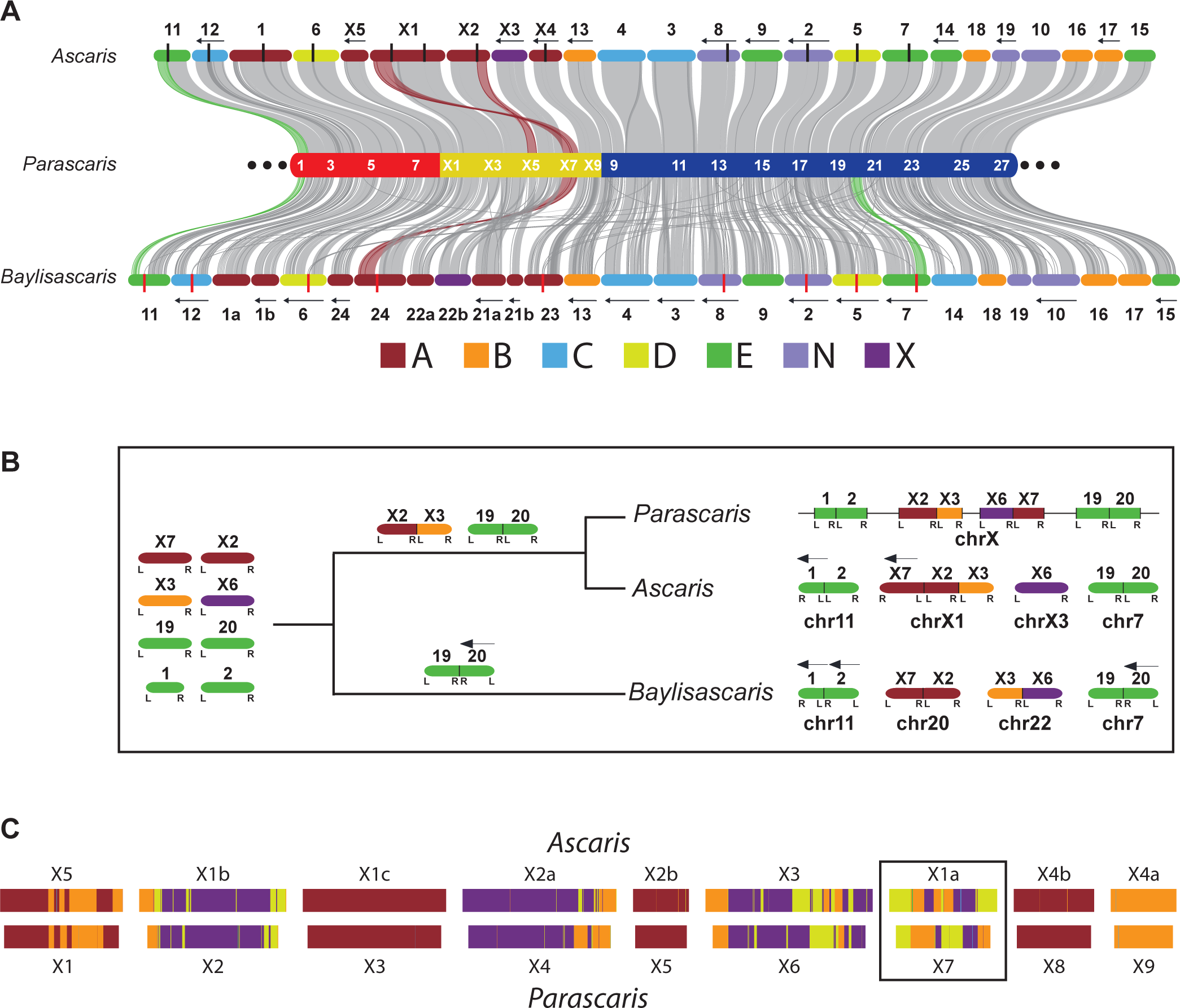
*Ascaris, Parascaris,* and *Baylisascaris* have the same somatic chromosomes. **A.** Synteny among three ascarid germline genomes. The *Parascaris* chromosome (middle) was colored as in Figure 1B, with the positions of every other somatic chromosome labeled in white. The *Ascaris* (top) and *Baylisascaris* (bottom) chromosomes were colored based on their predominant Nigon element (see legend for the seven Nigon groups). Their designated names for germline chromosomes are shown and their internal breaks for PDE are marked with black (*Ascaris*) or red (*Baylisascaris*) vertical lines. The left arrows on the top and bottom indicate chromosomes with reversed coordinates relative to *Parascaris*. See Figure S2 for synteny between *Ascaris* and *Baylisascaris*. **B.** An evolutionary model highlights various chromosome fusion events that lead to the different germline karyotypes observed in current ascarid genomes. All chromosomes were labeled on the top with designations from orthologous *Parascaris* somatic chromosomes; at the bottom are their current germline designations. The same Nigon color scheme was used. The L and R underneath each chromosome represent the left and right end of the ancestral chromosome, respectively. A chromosome with an R to L order indicates a reversed orientation relative to the *Parascaris* genome and is marked with a left arrow on the top of the chromosome. **C**. *Ascaris* and *Parascaris* sex chromosomes in the somatic cells were painted with the ancestral Nigon elements (see methods). The black box highlights the somatic chromosome X1a in *Ascaris* and X7 in *Parascaris* which have the most significant differences in their Nigon composition between the two species. See Fig S3 for highly conserved Nigon elements in the autosomes.

Analysis of the Nigon elements^20,41^ also revealed that each of the 36 somatic chromosomes in these ascarids originated predominately from a single ancestral Nigon group (Figure 4A and Table S3). One scenario for the origin of the current germline karyotypes is that they were derived from a single chromosome (as in *Parascaris*) that was split to become multiple germline chromosomes (as in *Ascaris* and *Baylisascaris*). This scenario is unlikely because the organization of the chromosomes is inconsistent among these ascarids (Figure 4A). Rather, our data points to a fusion model with the ancestor nematode having many smaller germline chromosomes (Figure 4B). Some ancient chromosomes in *Ascaris* and *Baylisascaris* and all ancient chromosomes in *Parascaris* have fused, with the fusion events happening at different times and/or with different chromosome ends or partners (Figure 4B). The fusion model is exemplified by the chr1/chr2 pair, where three of the four possible permutations of fusions were observed in the three nematodes (Figure 4B). The phylogenetic relationship of these nematodes also allows us to determine that the fusion of chrX2/chrX3 and chr19/chr20 pairs may have occurred before the divergence of *Ascaris* and *Parascaris* (Figure 4B). Overall, the data suggests ancient fusion events generated the different ascarid germline genomes, but PDE resolves these different fused germline chromosomes to a common set of 36 somatic chromosomes.

The organization of genes within the *Parascaris* sex chromosome region is more dynamic compared to the autosomes (Figures 4 and S2). To further elucidate the changes in the sex chromosomes between *Ascaris* and *Parascaris*, we used Nigon elements to paint the somatic sex chromosomes (and autosomes; see Figure 4C and Table S3). *Ascaris* somatic chromosomes are on average 13% larger than *Parascaris*, but the overall Nigon patterns along the orthologous chromosomes are consistent. Most sex chromosomes (and all autosomes) are painted with one dominant Nigon element. However, chromosomes X1a (*Ascaris*) and X7 (*Parascaris*) were composed of three Nigon groups (B, D, and X) that each occupy > 22% of the chromosome (Figure 4C). The order of the Nigon elements and the synteny (Figure 4A) between X1a and X7 reveal the chromosomes have undergone several inversions. Chromosome pair X3/X6 also showed inversions in their Nigon compositions and patterns (Figure 4C). The presence of multiple Nigon elements within these sex chromosomes is consistent with observations in other nematodes,^20,42,43^ suggesting that gene organization within the sex chromosomes can be more flexible compared to autosomes.

### Common features of nematode chromosomes indicate ascarid germline chromosomes were formed from fusions

The fusion model predicts that some remnant features of the pre-fused chromosomes may still exist in the current germline genomes. Typical features of a nematode chromosome include 1) a gene-rich region in the middle and repeat-rich region at the arms and ends;^34,44,45^ 2) more conserved and highly expressed genes in the middle;^34,44,45^ and 3) high recombination rates, SNP density, and GC content in the chromosome arms.^43,46-48^ We analyzed the ascarid genomes for signatures and/or relics of these common features in the presumptive 36 ancestral genomic regions (see Methods; Figure 5). We found that all 36 chromosomes show striking similarities in the distribution of GC content (Figures 5A and S3). They have elevated GC% at the arms and tips, and larger chromosomes often have high GC in the middle (see Figure S3) – an indication that these chromosomes could result from more ancient fusion events. Nevertheless, pairs of orthologous chromosomes between *Ascaris* and *Parascaris* mirror each other in their GC distributions (Figures 5A and S3). This data suggests that the GC profile is a stable feature of these 36 chromosomal regions; this profile likely has originated from ancient, separated chromosomes, supporting the fusion model.

**Figure 5.**
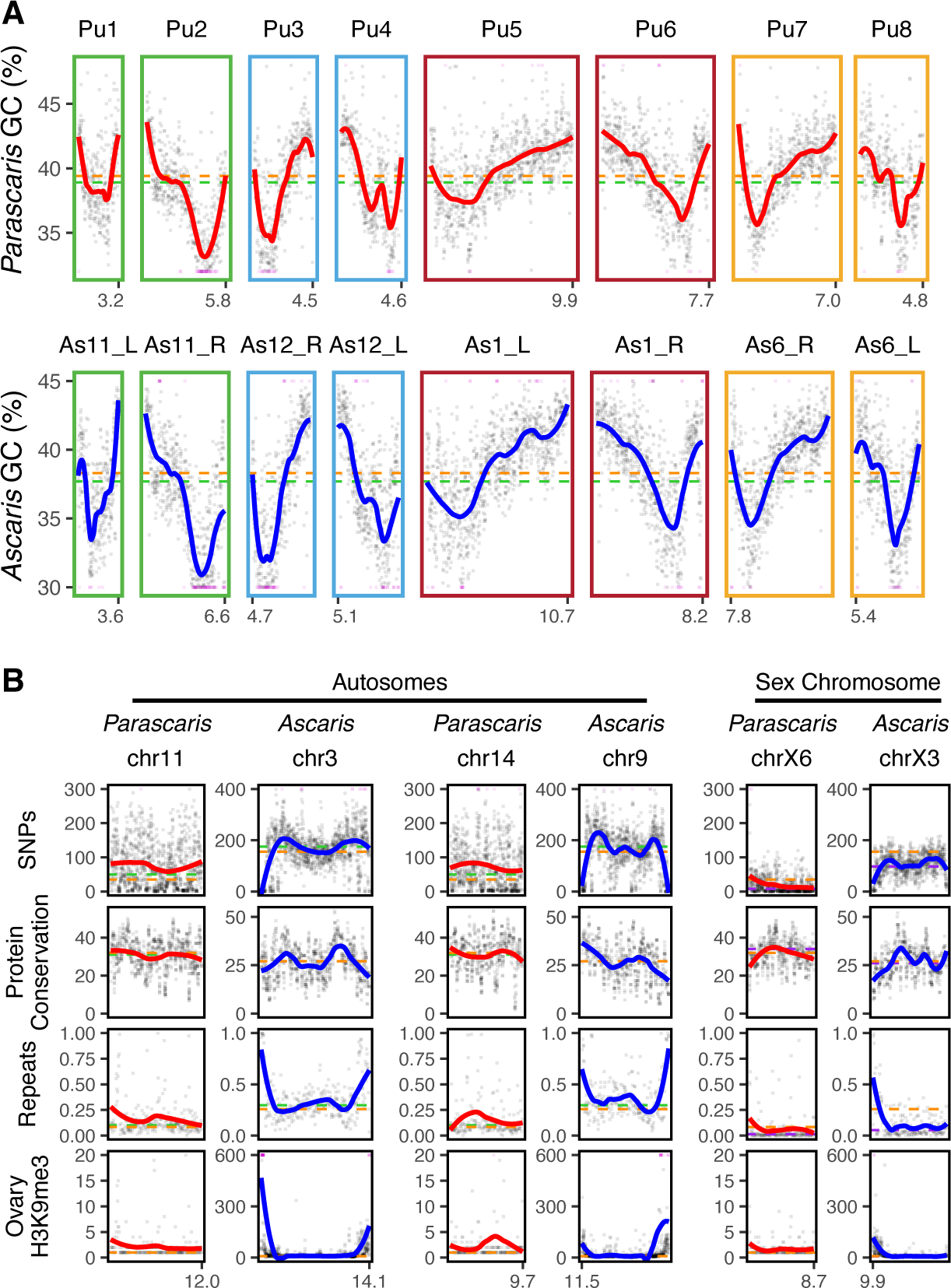
Common features of nematode chromosomes indicate fusions for ascarid germline chromosomes. **A.** GC content (%) was plotted for 8 presumptive ancestral pre-fused chromosome pairs in *Parascaris* and *Ascaris*. The ancestral Nigon group for each chromosome (see Figure 4A) is indicated by the color of the border. **B.** SNP density, protein conservation, repeat density, and H3K9me3 (from the ovary) were plotted in presumptive ancestral pre-fused chromosome pairs from *Parascaris* and *Ascaris*. Representative autosomes (left) and sex chromosomes (right) are shown. See Figure S3 for all 36 chromosomes. Each data point represents a binned region of the genome (window sizes for the bins: GC and SNPs = 10 kb; repeats and H3K9me3 = 50 kb; and protein conservation = 10 genes). A few data points beyond the y-axis ranges were converted to the minimum or maximum values and were colored in magenta. The colored trend lines (red or blue) were based on the LOESS regression. The horizontal dashed lines represent the average values of all chromosomes (orange), autosomes (green), or sex chromosomes (purple). The length of each chromosome is indicated at the bottom. Some *Ascaris* chromosomes are reversed in orientation (chromosome length labeled on the left side) to match their orthologous chromosomes in *Parascaris*.

Other features of the genome further support the germline fusion model. For example, the repeat density showed that the ends of the 36 presumptive chromosomes are generally enriched with repeats that are not the pentanucleotide and decanucleotide repeats (Figures 5B and S3). To further evaluate if these ends are enriched with heterochromatin, we determined the distribution of the histone mark H3K9me3 using CUT&RUN. Our data revealed that in the male and female germline tissues, H3K9me3 is highly enriched mostly at the ends of the 36 presumptive somatic chromosomes (Figure S3) and in the heterochromatic arms (Figure 3C). Notably, H3K9me3 is more enriched in autosomes than sex chromosomes (Figure S4C). In addition, our protein conservation analysis showed that proteins at the ends of chromosomes are less conserved (Figures 5B and S3). However, the SNP (and indel) density profiles between the *Ascaris* and *Parascaris* are not consistent (Figure 5B). The typical profile is maintained on the 24 *Ascaris* germline chromosomes rather than the 36 presumptive ancient chromosomes (Figure S5). In *Parascaris*, the SNP and indel distribution is more homogenous, with the sex chromosome region having a much lower level of SNPs and indels (Figures 5B, S3, and S4A-B). At the 240 Mb euchromatic chromosome scale, the density profiles also show higher SNPs and indels at the ends. These results are consistent with the observation in the nematode *Pristionchus* that chromosome fusions have repatterned the recombination rate and the SNP density.^47^ Thus, our analyses suggest that each of the 36 putative ancestral chromosomes has features that resemble a typical nematode chromosome. However, rapidly changing features, such as SNPs, indels, and recombination rates, are reflected in the fused germline genome. Together, our data on 1) the different sites and orders of fusions in karyotypes, 2) the independent Nigon blocks on the 36 somatic chromosomes, and 3) the common nematode chromosome features observed in the germline chromosomes support the model that ascarid germline genomes were fused from many smaller ancestral chromosomes.

### Dynamic organization of the *Parascaris* germline chromosome suggests a function for the heterochromatin arms during meiosis

The euchromatic region of the *Parascaris* represents only ∼10% of the germline genome. The rest of the genome is composed of tandem satellite repeats organized in the large heterochromatic arms (Figure 1A and Figure 6A). What is the function of these repeats in the germline and why do they need to be removed in the somatic cells? While we cannot fully assemble this genomic region or carry out genetic or functional analysis on these repeats, we used cytological analyses to characterize the organization of the arms and how they change during meiosis. The one-meter-long male germline reproductive tract allows easy separation of different stages of mitosis and meiosis (Figure 6B), enabling us to examine the dynamics of the chromosome throughout the mitotic divisions and spermatogenesis.

**Figure 6.**
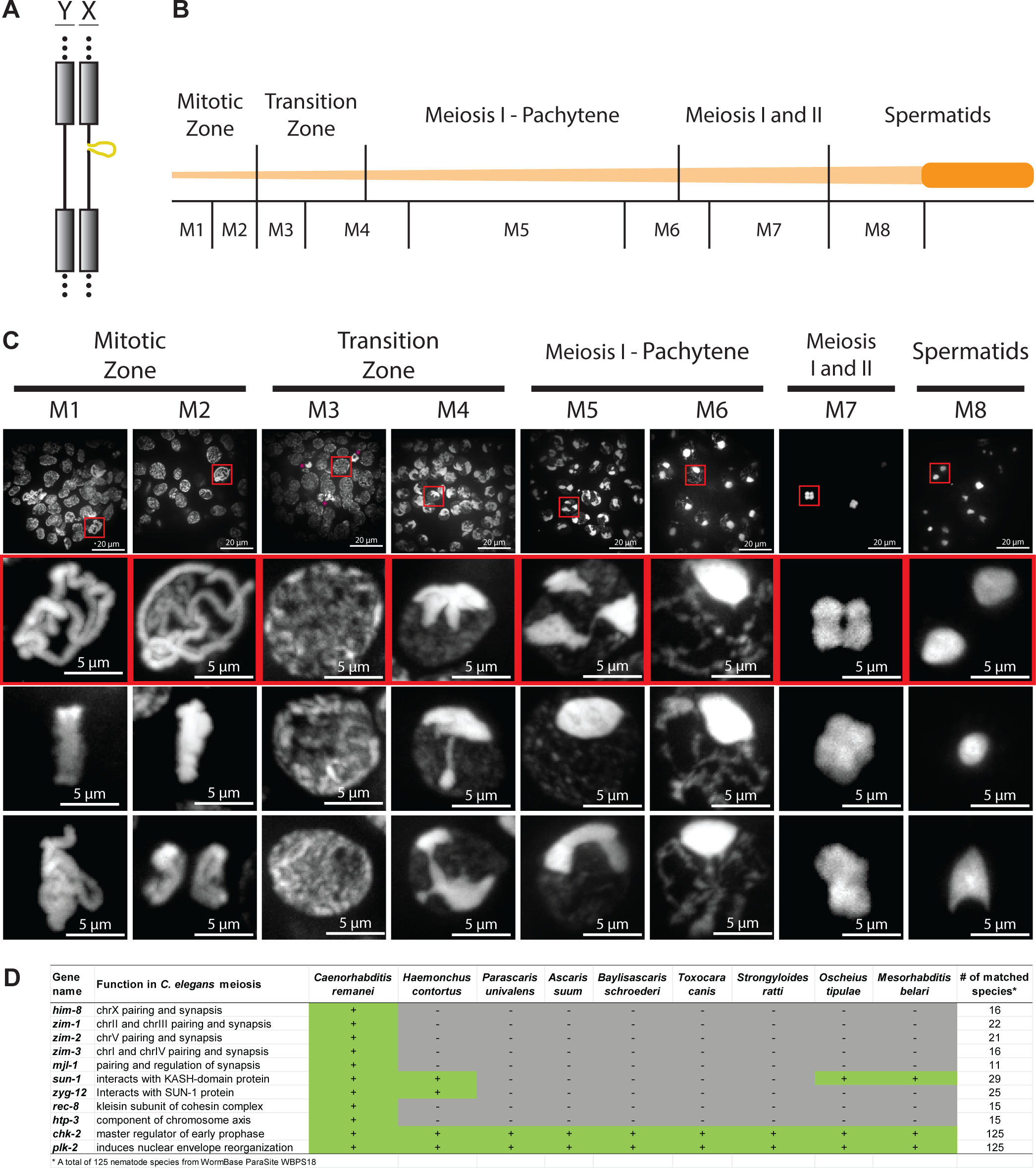
Dynamic organization of the *Parascaris* germline chromosome suggests a function for the heterochromatin arms during meiosis. **A.** An illustration of the X and Y chromosomes in the male germline. The X chromosome has a 55-Mb unique sex chromosome region (bulge) not found in the Y chromosome. The grey boxes and dotted lines represent the heterochromatic arms that will be eliminated by PDE. **B.** Schematic of the *Parascaris* testis showing different developmental stages (top) and regions sampled (bottom). A fully untangled *Parascaris* testis can reach one meter in length. **C.** Hoechst staining illustrates the dynamic organization of the germline chromosomes during male gametogenesis. The top row shows representative fields of view. The insets (red boxes) beneath show enlarged examples of condensed chromosomes at each stage. Additional images from the same stages were shown in the bottom two rows. Images were produced through maximum projection of multiple z-planes. See Movie S1 for the 3D organization of the chromosomes. **D.** Many genes involved in *C. elegans* pairing and synapsis are not found in nematodes with PDE. Known genes involved in *C. elegans* meiosis^49^ were searched against 125 nematode genomes available in WormBase ParaSite.^95^ Selected genes and species were listed, with genes *chk-2* and *plk-2* and species *C. remanei* and *H. contortus* serving as controls for detection. See Table S4 for the full list of known *C. elegans* genes involved in meiosis in all available nematodes.

We observed distinct features of the chromosome organization as the male germ cells undergo gametogenesis, as depicted as M1-M8 (Figure 6C and Movie S1). The heterochromatic arms and euchromatic regions are equally condensed in the mitotic (M1-M2) and early transition (M3) zones. The single pair of large, condensed chromosomes form a planar arrangement at the metaphase plate (see Movie S1). At the transition zone from mitosis to meiosis (M3 to M4), the presumptive heterochromatic repeats begin to differentially compress compared to euchromatin; they form various shapes as the heterochromatin congresses at one pole of the nucleus, reminiscent of the crescent-shaped chromatin polarized at the one side of the nuclei in *C. elegans*.^49^ The condensed heterochromatic repeats transition from multiple foci in early pachytene (M5) to a single focus at the nuclear periphery in late pachytene (M6), while the presumptive euchromatic regions form elongated, thread-like structure filling the rest of the nucleus, similar to the chromosome organizations observed in *C. elegans*^49^ and *Ascaris*.^50^ This organization brings together the germline chromosomes during pairing and synapsis and thus may function to better stabilize the structure of the chromosomes. The chromosomes further condense into a four-lobed metaphase structure in preparation for the first meiotic division (M7) and eventually become highly compact in spermatids (M8). The dramatic changes in the organization of the heterochromatic arms during *Parascaris* gametogenesis, particularly the structural changes during pachytene (M5-M6), suggest a potential function for these chromosome arms during meiotic pairing and synapsis.

In most nematodes, homologous chromosomes are paired during meiosis to allow synapsis, recombination, and crossover.^49,51,52^ However, in some unichromosomal asexual nematodes such as *Diploscapter pachys*, the single pair of chromosomes does not undergo pairing and crossover, resulting in significant heterozygosity in the two chromosomes.^53^ *Parascaris* is also unichromosomal, but it undergoes sexual reproduction and has a low level of heterozygosity in the euchromatic region, suggesting paring and crossover occur during meiosis. Comparative genomics indicates many genes involved in *C. elegans* meiotic pairing^54-57^ are missing in *Parascaris* and other nematodes with PDE (Figure 6D and Table S4). Although it is possible these genes are too diverged to be identified through sequence homology, it seems more likely that *Parascaris* and other PDE nematodes use different mechanisms for pairing.^52,58^ During pairing and synapsis, the lack of the 55 Mb sex chromosome region in the Y chromosome may also cause instability in the formation of the synaptonemal complex (Figure 6A). We reason that the anchoring of the large heterochromatic arms at the nuclear membrane may stabilize the structure of the synaptonemal complex and thus facilitate pairing (Figure 6C). Overall, the dynamic reorganization of heterochromatic arms with their high level of compaction in pachytene suggests they could serve a function for pairing and stabilizing the chromosome structures during meiosis.

## Discussion

*Parascaris* served as a classical model organism for early studies on chromosome biology.^23,24^ Here, we assembled the euchromatic region of the single germline chromosome and analyzed sequence and karyotype changes that occur during *Parascaris* programmed DNA elimination. We also compared the PDE features and karyotypes to other related ascarids. Our results prompt a model where the fusion of ancient chromosomes in these nematodes formed their germline chromosomes and these fused chromosomes are subsequently resolved in the somatic cells by PDE, suggesting PDE is a part of the mechanisms involved in the evolution of these nematode genomes.

Many metazoans have been identified to undergo PDE, but the functions of PDE are largely unknown due to the lack of functional and genetic models.^8,19^ Genomic and cytology data suggest several potential functions for nematode PDE: 1) silencing of repeats and genes associated with germline development through their elimination in the somatic cells;^2,10^ 2) remodeling of chromosome ends in the somatic cells;^14^ 3) changing chromosome karyotypes;^14^ 4) facilitating the evolution of novel genes;^2,8^ and 5) sex determination (limited to *Strongyloides*).^9^ Our data in *Parascaris* supports most of these functions and highlights PDE’s role in karyotype changes between the germline and the somatic cells. *Parascaris* provides an ultimate model for the fusion and fission of chromosomes, with a single germline chromosome resolved into 27 autosomes and 9 sex chromosomes and a switch from an XY sex determination karyotype in the germline to an XO karyotype in the somatic cells.

Fusion and fission are major mechanisms that lead to changes in chromosome karyotype during evolution.^20^ In nematodes, these could in theory be more prevalent due to the holocentric nature of the chromosomes.^32-34^ Our analyses of the ascarid chromosomes and PDE lead to a model of chromosome evolution in nematodes, as exemplified in the sex chromosome regions (Figure 7). We believe that many smaller chromosomes existed in the ascarid ancestral germline genome. These ancestral chromosomes would have undergone PDE – since if no PDE has occurred, a single DNA break at a fusion site would be sufficient to break the fused chromosome and there would be no loss of DNA. However, at each presumptive fusion site, two breaks are always observed flanking an extension of eliminated DNA (Figure 2A), suggesting each break likely originated from the end of an ancestral chromosome that underwent PDE with DNA loss. Various chromosome fusion events occurred during evolution leading to the current different organizations of the germline chromosomes in ascarids (Figure 7). In addition, the resulting differences in karyotypes could cause incompatibility in homologous chromosome pairing, suppression of recombination, and reproductive barriers, and consequently lead to speciation.^59^ The inherited PDE mechanism is used to split the germline chromosomes into the ancestral, pre-fused somatic chromosomes. Since the PDE mechanism is precise, it allows a reproducible regeneration of the smaller ancestral chromosomes with their ends removed (Figure 7). This model suggests that PDE is an ancient mechanism in nematodes that existed in the common ancestor of ascarids before the germline chromosome fusion events; it serves as a mechanism for chromosome fission and is part of the evolutionary repertoire that shapes nematode genomes.

**Figure 7.**
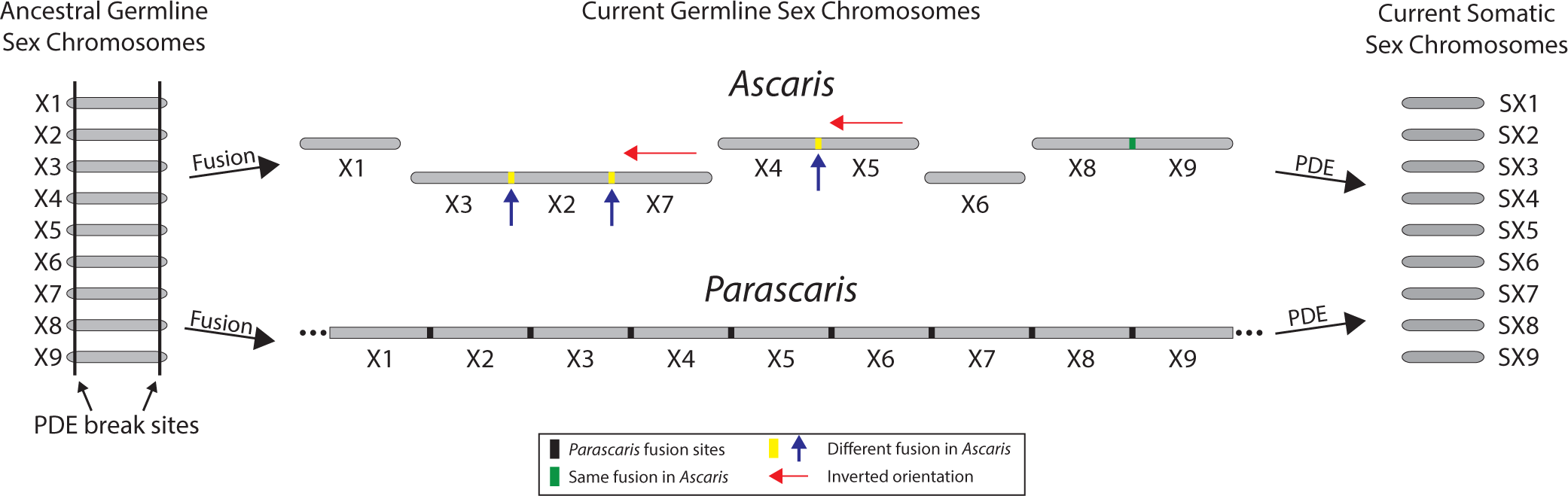
Fusion, PDE, and evolution of the sex chromosomes in parasitic nematodes. A model showing the chromosome fusions and their split by PDE in sex chromosomes. In the ancestral state, all chromosomes likely undergo PDE at both ends of the chromosome (left). Chromosome fusion events lead to different germline chromosomes (and likely speciation) in nematodes. See the legend for the different types of fusion sites and chromosome orientations (middle). PDE splits different germline chromosomes into the same somatic chromosomes and restores the pre-fused germline karyotype (right). Autosomes, not shown in this model for simplicity, follow this same evolutionary trajectory and fate as the sex chromosomes.

What is the advantage of a fused nematode chromosome in the germline and why does it need to be split in the somatic cells? During meiosis, homologous chromosomes need to be paired and faithfully segregated;^51,52,58^ mis-segregation can lead to aneuploidy and other chromosome abnormalities.^60^ Maintaining many chromosomes requires a pairing system^58^ that can accurately and efficiently identify all pairs of homologous chromosomes. Thus, reducing the number of germline chromosomes through fusion could alleviate the burden of the pairing system and decrease nondisjunction errors caused by mis-segregation. The need for accuracy and efficiency in meiosis is consistent with the fecundity of these parasitic nematodes that produce hundreds of thousands of viable eggs per day.^11^ In addition, nematode chromosome fusion could also repattern recombination rates and facilitate reproductive isolation, promoting evolution and speciation.^47^ However, fused nematode chromosomes may lead to 3D genome organization changes that are important for gene expression and regulation during development.^61^ A typical nematode chromosome has a gene-rich region in the middle and repeat-rich areas at the arms and ends (see Figure 5). This organization is conserved suggesting its biological significance. It could be responsible for more effective transcription, silencing via heterochromatin formation, and gene regulation.^62,63^ We reason a fused chromosome in the somatic cells may disrupt these functions and regulation. Therefore, fusion and PDE together may provide a solution for organisms to benefit from fewer chromosomes in the germline and the typical organization of nematode chromosomes in the somatic cells.

In *C. elegans*, the pairing of homologous chromosomes is through the pairing centers residing at one end of each chromosome, with four zinc-finger proteins recognizing six chromosomes.^52,54-57^ Interestingly, we did not find homologs of these proteins in ascarids and many other nematodes (Figure 6D), suggesting some nematodes may use different pairing mechanisms. The need for meiotic pairing but the apparent lack of a conserved pathway have created opportunities for diverse pairing mechanisms to evolve and be adapted in nematodes.^58,64^ A plausible candidate for an alternative pairing mechanism is the repetitive sequences in the subtelomeric regions and the telomeres.^52,58^ In many organisms, the formation of a telomere bouquet appears to facilitate the recognition of homologs by limiting the chromosomes in a concentrated area for pairing.^58,65,66^ Furthermore, it was demonstrated in wheat that the subtelomeric regions are involved in the correct pairing of homologous chromosomes.^67^ Future studies on the pairing mechanisms are warranted to elucidate their variations in diverse nematodes.

The heterochromatic arms in *Parascaris* (and the 121-bp satellite repeats in *Ascaris*^8^) may further facilitate pairing by stabilizing the chromosomes during meiosis (Figure 6D and Movie S1). However, in the somatic cells of PDE species, the sequences (the germline telomeres and repeats) for pairing are no longer needed and their pairing capability could pose a hindrance to mitosis and/or genome organization. Our fusion and fission model suggests that the internal breaks can be viewed as breaks at the ends of ancient chromosomes. This is consistent with the model that PDE removes the pairing centers at the ends of the germline chromosomes. In addition, the large amount of heterochromatic DNA (90%; 2.2 Gb) in *Parascaris* is not economical to replicate and maintain. PDE removes these repeats and telomeres thus streamlining the genome in the somatic cells. Consistent with this, we found that after PDE in the nematode *O. tipulae*, the length of new telomeres in the somatic cells is often much shorter than the germline.^19^ Overall, we speculate that PDE may serve a function to discard DNA that is nonessential in the somatic cells but has an important role in the pairing of homologous chromosomes during germline meiosis.

## Conclusions

PDE can result in dramatic genome changes, yet the biological significance of this process has remained obscure in most PDE models. We demonstrated that through the fusion and fission of the chromosomes, PDE could be a mechanism that facilitates genome maintenance. These seemingly paradoxical aspects of PDE highlight the dynamic nature of the chromosomes and how evolution adapts and uses mechanisms that modify chromosomes to maintain the genome. The phenomenon of PDE has been recognized for over a century, yet we still know little about the function(s), molecular mechanism(s), and evolution of PDE in multicellular organisms. Future studies on the growing list of metazoans with PDE, many amenable to genetic manipulations,^68^ are promising to provide novel insights specifically into PDE and more broadly into chromosome dynamics.

## METHODS

### Lead Contact

Further information and requests for resources and reagents should be directed to and will be fulfilled by the Lead Contact Jianbin Wang.

## Materials Availability

Materials from this study are available on request.

## Data and Code Availability

All sequencing data were deposited at the NCBI SRA (accession number: pending) and GEO (accession number: pending) databases. The data for genome sequencing, gene models, RNA-seq, and CUT&RUN are also available in a UCSC Genome Browser track data hubs that can be accessed with this link: https://genome.ucsc.edu/s/jianbinwang/parascaris. In addition, gene models, annotation, and expression datasets are available at https://dnaelimination.utk.edu/protocols-data/.

## Method Details

### *Parascaris* material and high-molecular-weight DNA isolation

Collection of *P. univalens* tissues, zygotes, and embryonation were as previously described.^13^ Dissected testis, ovary, and intestine were frozen in liquid nitrogen and stored at -80°C; fertilized eggs were stored at 4°C. DNA isolation for PacBio sequencing was prepared from testis using phenol/chloroform extraction. Briefly, the testis tissue was ground into a fine powder with liquid nitrogen using a mortar and pestle and was digested overnight at 55 °C with 0.5% SDS and 150 μg/mL proteinase K (Ambion, AM2548) in buffer (50 mM Tris-HCl pH 7.4, 100 mM EDTA, and 100 mM NaCl). The DNA was extracted twice with phenol/chloroform (1:1) and once with chloroform/isoamyl alcohol (24:1), followed by ethanol precipitation using 0.3 M NaOAc pH 5.2, and 2.5 volumes of 100% ethanol. The precipitated DNA was dissolved in low TE buffer (10 mM Tris-HCl pH 8.0, 0.1 mM EDTA) or water.

### PacBio sequencing and initial assembly

A PacBio CLR library derived from testis DNA was prepared and sequenced by the University of Washington PacBio Sequencing Services (https://pacbio.gs.washington.edu/) using the Sequel II system. From one SMRT Cell run, we obtained ∼172 Gb (∼69X) raw reads with an average read length of 14 kb. Reads (∼60% of total bases) that are composed largely of satellite repeats (5-mer and 10-mer) were identified (using cross_match) and filtered out. The remaining reads (4,046,075 reads, 67.9 Gb with an average read length of 16.8 kb) were error-corrected with Canu (correctedErrorRate=0.105) to yield 3,662,170 corrected reads. Corrected PacBio reads over 20 kb (average 30 kb, total 34.4 Gb) were used to assemble the initial germline genome assembly using Canu^69^ with parameters “corOutCoverage=1000 correctedErrorRate=0.105 corMinCoverage=0 genomeSize=280m”. The initial assembly had 1,151 contigs with N50 = 2.08 Mb (N50 number = 46).

### *Parascaris* nuclei isolation and Hi-C library preparation

Samples from both germline (testis and ovary) and somatic cells (intestine) were fixed according to the Arima Hi-C 2.0 protocol for standard input. To collect nuclei for Hi-C, samples were ground into a fine powder under liquid nitrogen and were fixed in 5.5 mL of fixative solution (1X PBS, 0.4 mM PMSF, 10 mM NaCl, 0.5 mM EDTA, 0.25 mM EGTA, 25 mM HEPES pH 8.0, 2.2% formaldehyde) while rotating at room temperature for 20 minutes. A 289-µL stop solution was added and rotated at room temperature for 10 minutes. Samples were spun down at 2,500 x g for 5 minutes, washed with 1 mL 1X PBS, 0.4 mM PMSF, aliquoted, pelleted, and the supernatant removed, and stored at -20°C.

Hi-C was performed using the Arima-HiC 2.0 kit according to the manufacturer’s protocol (Arima, Cat#A410110). Following Hi-C, samples were fragmented using a Covaris M220 ultrasonicator (Covaris, Cat#500295) to yield fragments of around 500 bp long. Size selection was performed using AMPure XP magnetic beads (Beckman Coulter, A63881). Libraries were constructed for Illumina sequencing using the Swift Biosciences Accel-NGS 2S Plus DNA Library kit (Swift Cat# 21024) and the KAPA Library Amplification kit (Roche Cat# KK2620) according to the manufacturer’s protocols. Sample concentrations were determined using a DNA Qubit (Invitrogen, Cat# Q33238), and library fragment sizes were analyzed using a TapeStation 4200 (Agilent, G2291BA). These and all subsequent Illumina libraries were sequenced using a Novaseq 6000 at the University of Colorado Anschutz Medical Campus Genomics Core.

### Genome scaffolding

PacBio sequencing and Hi-C data were used to scaffold contigs into a single chromosome-level assembly for the germline chromosome in *Parascaris*. The 1,151 contigs from the Canu assembly were first scaffolded into 896 fragments using SALSA.^70^ These scaffolds were further analyzed for interactions between their ends using Hi-C data through several iterations of scaffolding. The assemblies produced through each round were analyzed using HiC-Pro^71^ and Juicebox^72^ to guide the scaffolding process. The final assembly is composed of one large scaffold that includes the euchromatic region of the single germline chromosome (size 240 Mb) and 50 small unplaced contigs in 3.6 Mb (1.5%) that are enriched with repetitive sequences.

### RNA-seq

We generated 12 new RNA-seq libraries using methods previously described.^19^ Briefly, total RNA was prepared using the TRIzol (Invitrogen Cat# 15596026) protocol. The quality of the RNA was evaluated with the TapeStation 4200 (Agilent) and quantified with the Qubit4 Fluorometer (Thermo Fisher). The ribosomal RNAs were removed using the RiboCop rRNA Depletion Kit (LEXOGEN Cat# 144). The RNA-seq libraries were constructed using the CORALL Total RNASeq Library Prep Kit (LEXOGEN Cat# 146).

### Gene models and annotation

We used 25 RNA-seq libraries (13 were from a previous study^13^) that cover 17 major stages of nematode development (see Table S1) to identify genes in the *Parascaris* genome. We first removed ribosomal RNA reads using bowtie2,^73^ then mapped the non-rRNA reads against the *Parascaris* genome using STAR.^74^ Transcript assembly was performed using the StringTie^75^ to generate a non-redundant set of transcripts. Candidate coding regions were determined using TransDecoder (https://github.com/TransDecoder). Cryptic transcripts, artifacts, and transcriptional noises were further removed using the established criteria in a previous study.^19^ Overall, we identified 16,603 protein-coding genes and 56,514 RNA transcripts. The genes were used to BLAST^76^ against protein databases (NCBI nr, UniProt,^77^ Swiss-Prot, and WormBase^78^) to assign annotation (Tables S1).

### Genes expression analysis

HTSeq^79^ was used to count the number of reads for each transcript. To identify differentially expressed genes between the developmental stages, we used the DESeq2^80^ package with an adjusted p-value of less than 0.05 and a fold-change cutoff of 2. To gauge the tissue-specific expression of the genes, we used the average value of RNA-seq data from 8 broadly defined tissues from *Parascaris*, including testis mitosis (M1-M3), testis meiosis (M5-M7), ovary, early embryos (1-32 cells), late embryos (>64 cells), carcass, intestine, and juvenile (mainly composed of somatic cells). The tissue-specific expression was scored as described.^14^ Briefly, the RNA levels (rpkm) for each gene were compared among all 8 tissues. A gene is considered highly tissue-specific (score = 100) if its expression is 4-fold higher in one tissue than any of the other 7 tissues. The fewer the number of tissues that meet the 4x expression cutoff the less tissue-specific expression the gene is. Genes with a maximal expression rpkm <= 5 in all 8 tissues were considered as low expressed genes and were excluded from the tissue-specific expression analysis.

### CUT&RUN and data analysis

*Parascaris* tissues were ground to a fine powder using a mortar and pestle with liquid nitrogen. Each CUT&RUN reaction used approximately 50μl of tissue powder. CUT&RUN was done using the EpiCypher CUTANA ChIC-CUT&RUN kit (SKU: 14-1048). For each reaction, 0.5μg of antibody against H3K9me3 (CMA318), H3K4me3 (CMA304), or CENP-A (#325 and #345)^15^ was used. Sequencing library preparation was done using a 2S Plus DNA Library Kit from Swift/IDT (Swift Cat# 21024). CUT&RUN reads were mapped to the *Parascaris* genome using bowtie2^73^ and processed using SAMtools^81^ and BEDTools.^82^ The reads were normalized based on the genome-mapped reads. CENP-A CUT&RUN data were analyzed using the same method described in *Ascaris* CENP-A^15^ to define the CENP-A enriched domains. CUT&RUN data was uploaded to the UCSC genome browser track data hubs.^83^

### Repetitive sequence identification

Repetitive sequences were identified using a combination of homology-based and *de novo* approaches, including RepeatMasker,^84^ LTRharvest,^85^ RepeatScout,^86^ RepeatExplorer,^87^ and dnaPipeTE.^88^ The predicted repeats were filtered for redundancy with CD-hit.

### Gene prediction in *Baylisascaris*

*Baylisascaris* RNA-seq reads were mapped to the *B. schroederi* genome using STAR to identify genes as described above. To uncover potential genes missed due to the limited developmental stages sampled from the available RNA-seq libraries (such as testis-specific genes), we also used AUGUSTUS^89^ to predict genes in the *B. schroederi* genome. First, 200 training gene structures from the *B. schroederi* genome and protein sequences were generated using GenomeThreader (https://genomethreader.org/). The resulting gtf training gene file and the genome file were converted to GenBank format and used to train AUGUSTUS. Predicted genes were merged with the RNA-seq-based gene models to produce the final gene models for comparative analysis.

### Genome synteny and Nigon element analysis

Pairwise comparisons among *Ascaris, Parascaris,* and *Baylisascaris* were carried out to obtain the reciprocal best blast hits for orthologous among these nematodes. Syntenic blocks were identified using the coordinates of the genes along the assembled chromosomes. To determine the presumptive fusion sites, orthologous regions (tBLASTx) were used to define the boundary of ancestral chromosomes using the germline chromosome ends of the *Ascaris* genome. However, for fused chromosomes that have the same orientation in both *Ascaris* and *Parascaris* germline genomes, the middle of the eliminated regions were chosen as the presumptive fusion sites. Plots were generated using Circos,^90^ Synvisio,^91^ and R packages (tidyverse, readr, scales, and patchwork). For Nigon element analysis, the set of predicted genes in the *Parascaris* genome was searched for orthologs to a set of 2,191 *C. elegans* genes defined as Nigon elements^20^ using BLAST.

### SNP and indel analysis

Illumina sequencing reads were first mapped to *Parascaris* genome assembly. Short variants (SNPs and indels) were then called using the genome analysis toolkit (GATK) and the HaplotypeCaller package^92^ with default parameters. SNP and indel density were calculated per megabase.

### *Parascaris* immunohistochemistry on CENP-A

The staining procedure was adapted from a previous study on *Ascaris* embryos.^93^ Briefly, embryos were fixed in 500 uL of a fixative solution containing 50% methanol (Fisher Chemical, A412SK-4) and 2% paraformaldehyde (Ted Pella, Inc., Cat # 18505) in 1X PBS buffer. The embryos were freeze-cracked at least five times and rehydrated by incubation in 25% methanol in 1X PBS, followed by incubation in 1X PBS. Embryos were resuspended in 1 mL of blocking solution (0.5% BSA in 1X PBS) and incubated for 30 minutes at room temperature with rotation, then they were put in blocking solution and stained with primary antibody (diluted 1:1000) overnight at 4°C with rotation. Secondary antibody incubation (diluted 1:1000) was carried out at room temperature for two hours in the dark with rotation. Embryos were then incubated with 500 µL of Hoechst 33342 (1 µg/mL) (Invitrogen, Fisher Cat# H3570) dye at room temperature for ten minutes in the dark with rotation. Embryos were washed in a blocking solution and mixed with Vectashield plus antifade mounting media (Vector Laboratories, H-1900-10). Imaging was performed on a Nikon Eclipse Ti inverted spinning disk confocal microscope using a 100X 1.49 NA oil immersion objective lens. Samples were mounted to slides using Vectashield Plus antifade mounting media (Vector Laboratories, H-1900-10) and imaged using a Nikon Eclipse microscope equipped with a Yokogawa CSU22 scanning laser confocal unit, a Photometrics Evolve 512 emCCD camera, and a 100X, 1.49 NA lens. Image manipulation was performed in FIJI.^94^ All contrast adjustments in figures are linear.

### *Parascaris* male germline microscopy

Intact testes were dissected from adult male worms and fixed (long-term) in 2% paraformaldehyde. Tissues were cut into small pieces approximately 2.5 cm long at developmental regions defined by relative distance to the distal tip. Germline pieces were placed in a drop of 1X PBS buffer pH7.4 on a glass slide. A glass rod was rolled over the tissue to extrude germline cells from the somatic sheath surrounding them. Samples were stained in Hoechst-33342 (Fisher Scientific, H3570) at room temperature for ten minutes and were rinsed in 1X PBS buffer. Microscopy and imaging analysis were done as described in the immunohistochemistry section.

## Supplemental information

### Supplemental figures

**Figure S1.**
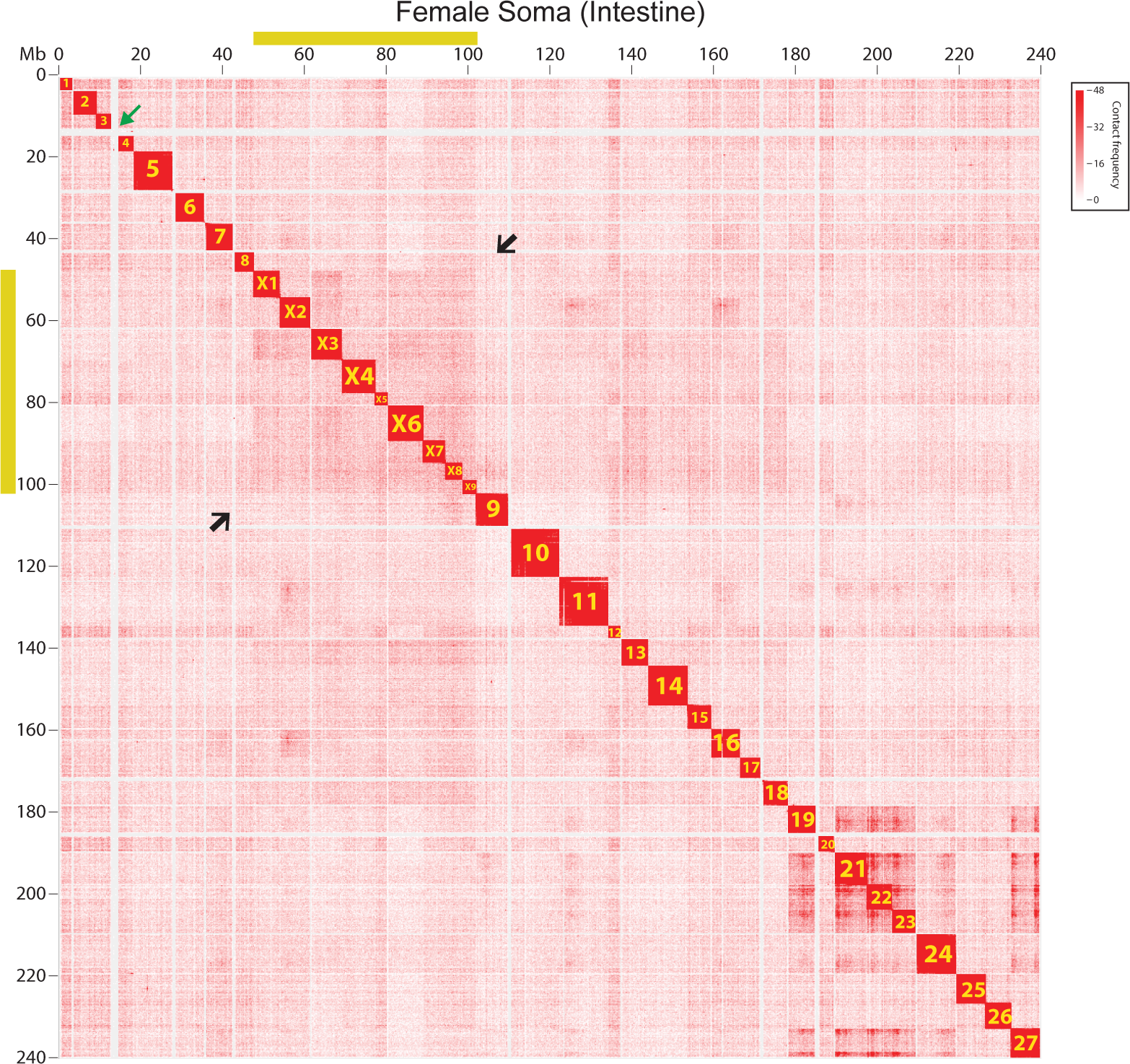
*Parascaris* Hi-C interaction heatmap for female intestine. Yellow bars along the x and y axes indicate the sex chromosome region of the *Parascaris* X chromosome. The black arrows point to the junctions between autosome and sex chromosome regions and the green arrow shows the largest piece of eliminated DNA.

**Figure S2.**
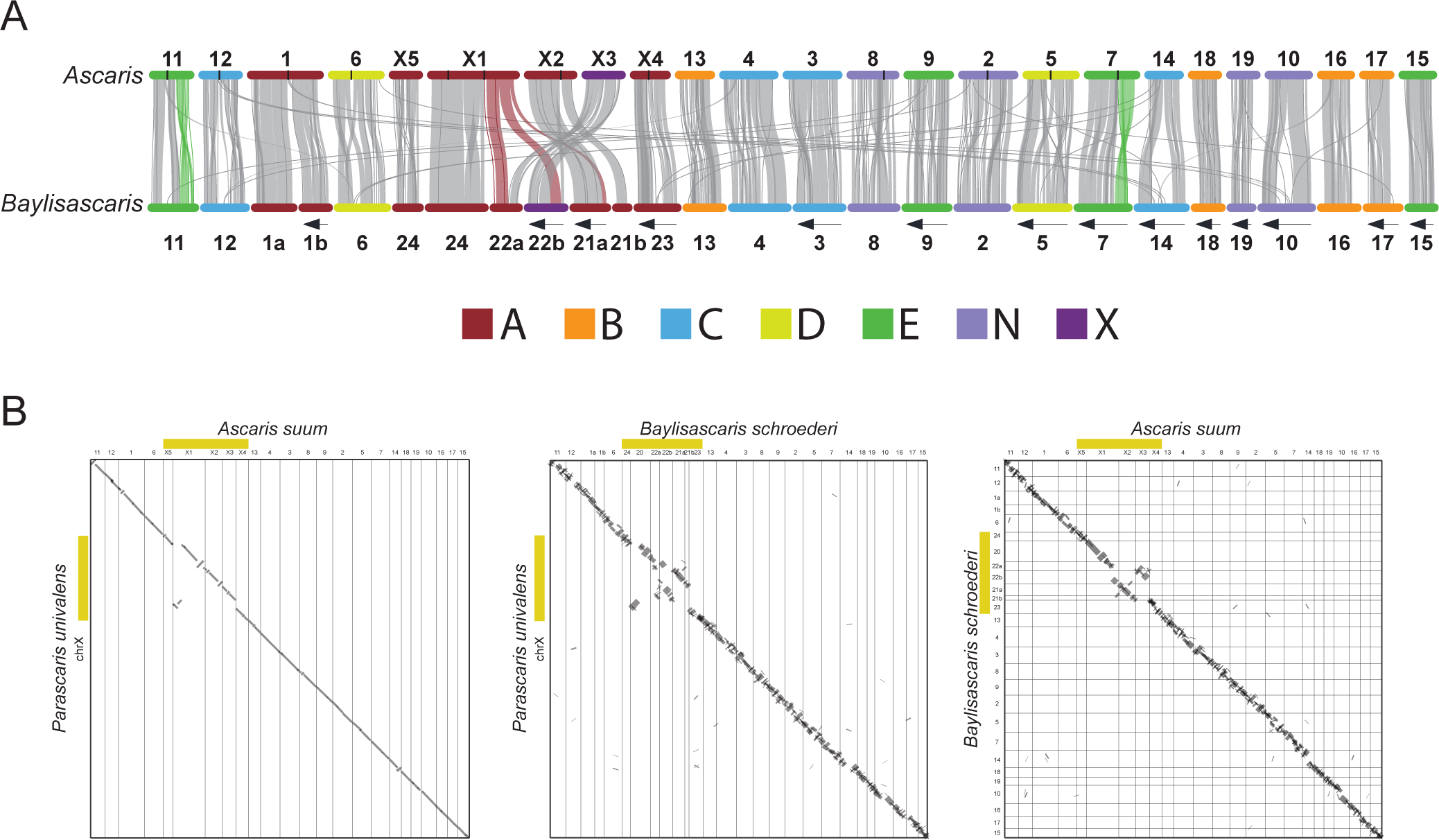
Synteny among *Ascaris, Baylisascaris* and *Parascaris*. **A.** The *Ascaris* and *Baylisascaris* chromosomes were colored based on their predominant Nigon element (see legend for the seven Nigon groups). Their designated names for germline chromosomes are shown and their internal breaks for PDE are marked with black (*Ascaris*) or red (*Baylisascaris*) vertical lines. The left arrows on the top and bottom indicate chromosomes with reversed coordinates. **B.** Pairwise comparisons between ascarids germline genomes in dot plots. Sex chromosome regions were indicated in yellow bars.

**Figure S3.**
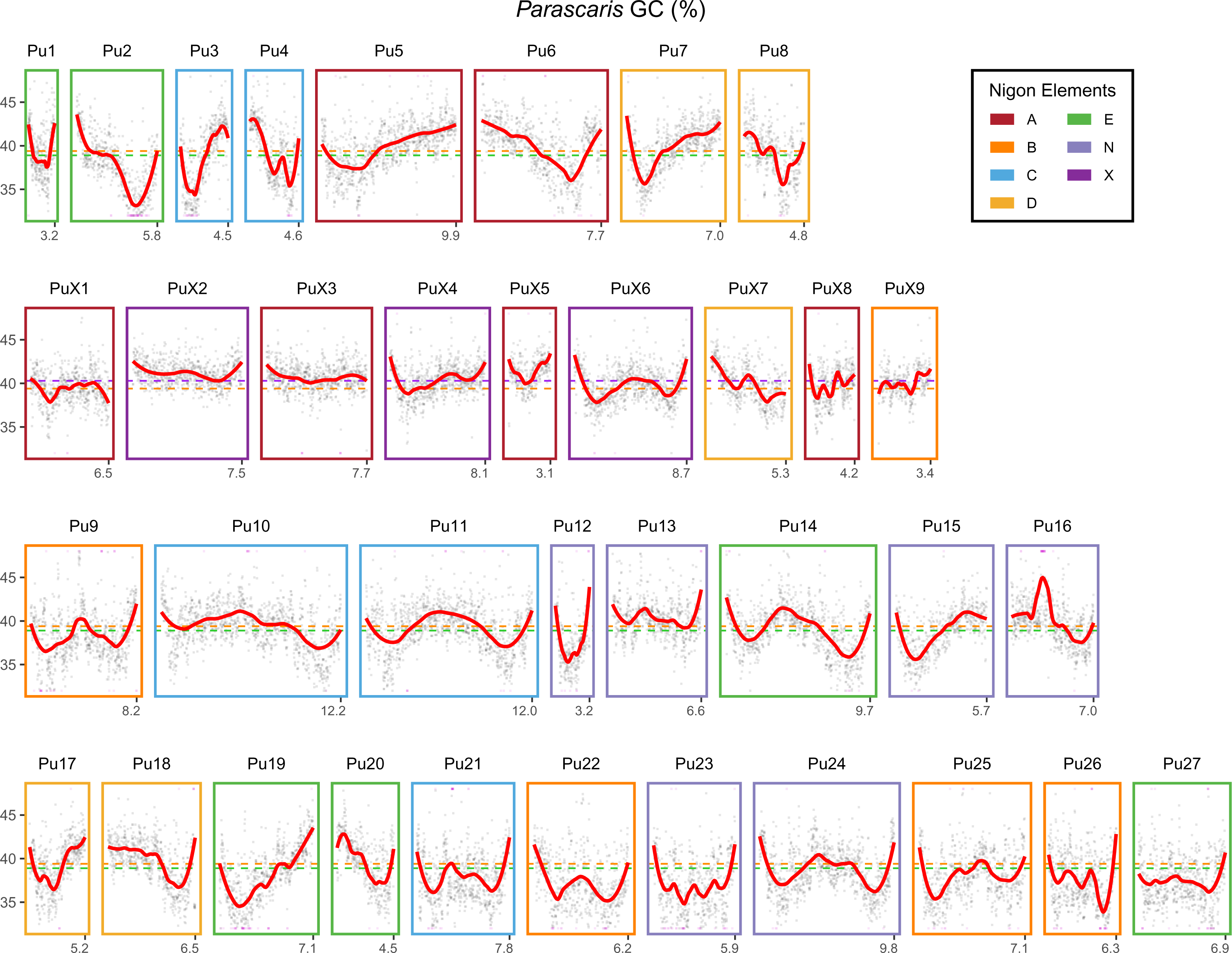
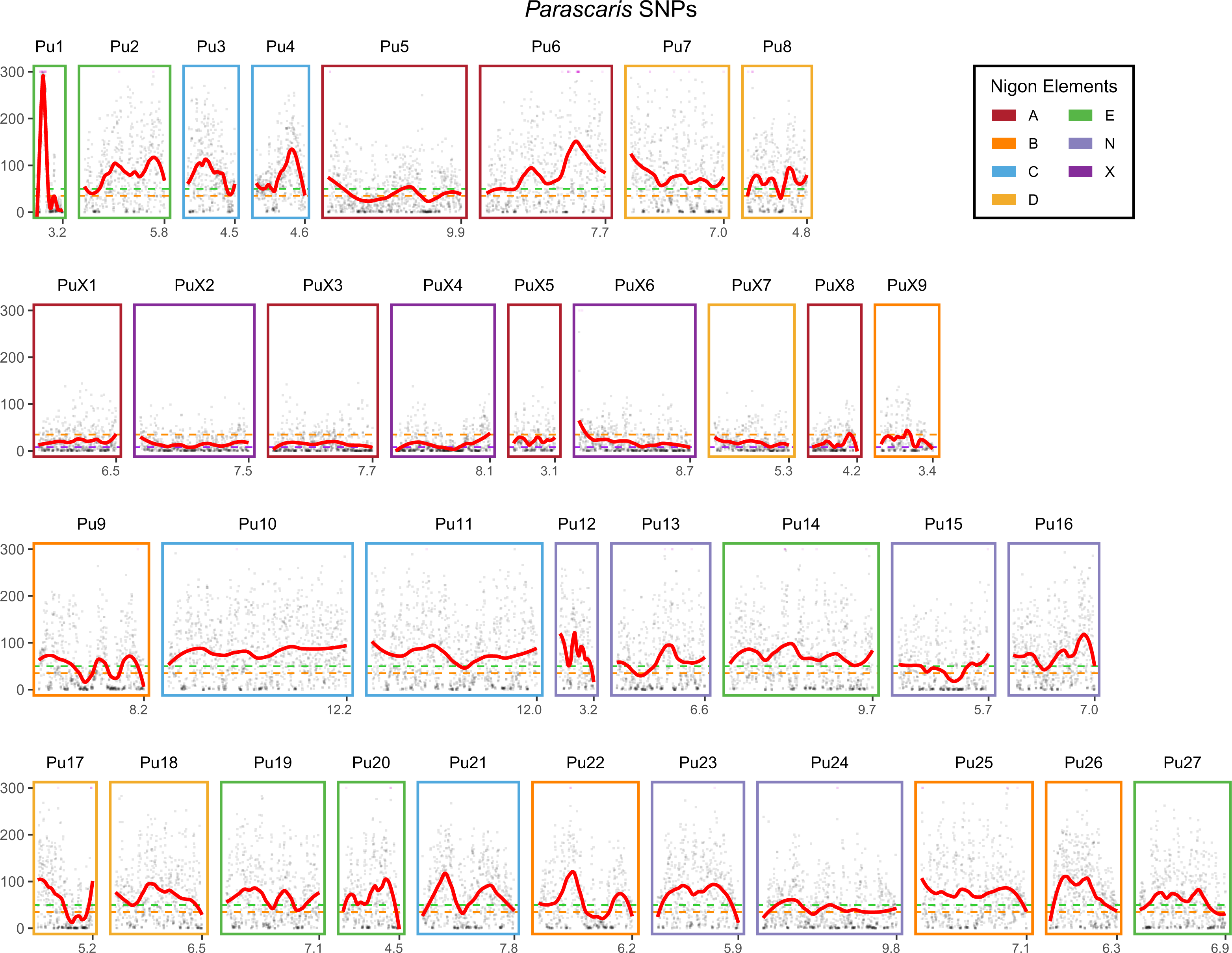
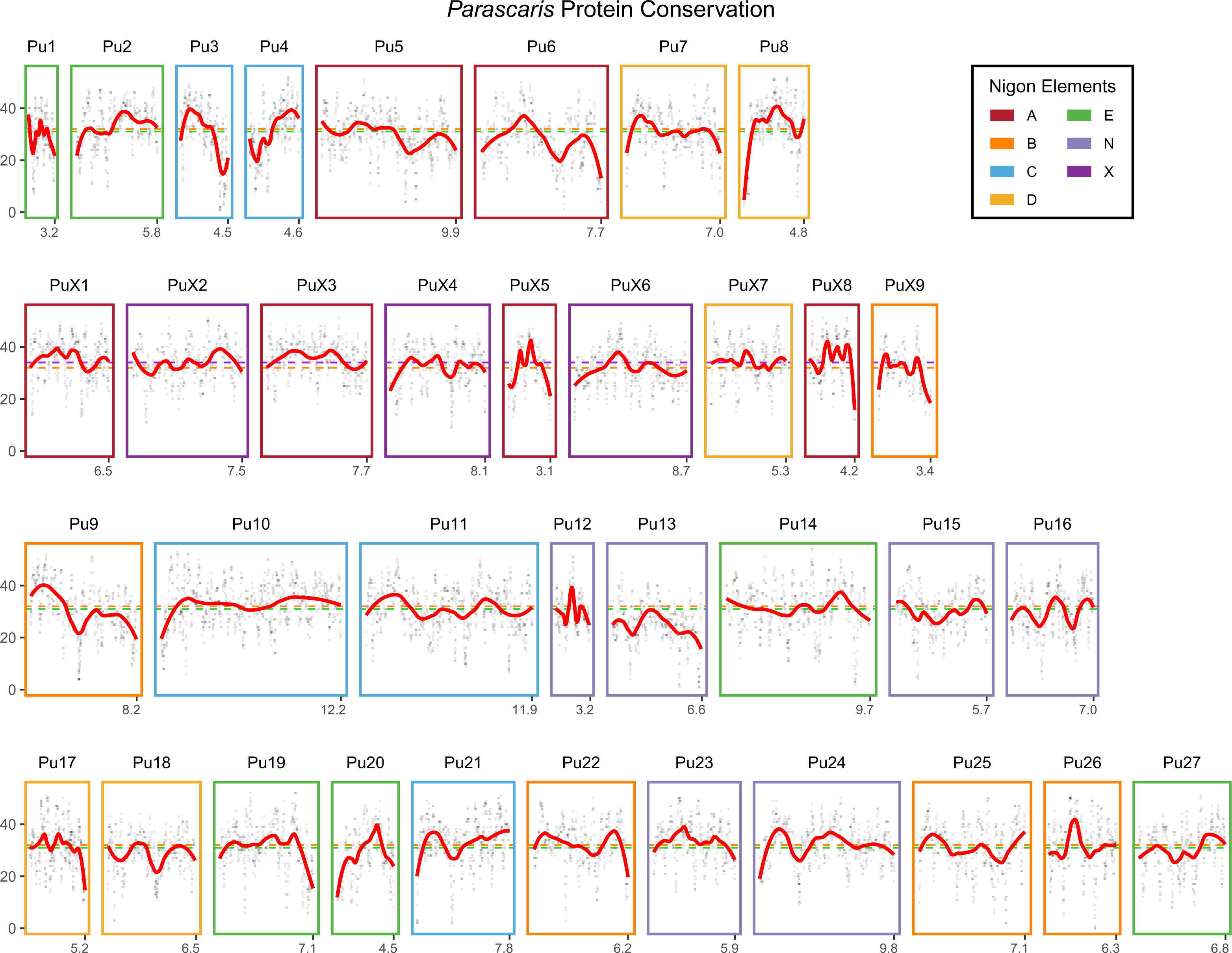
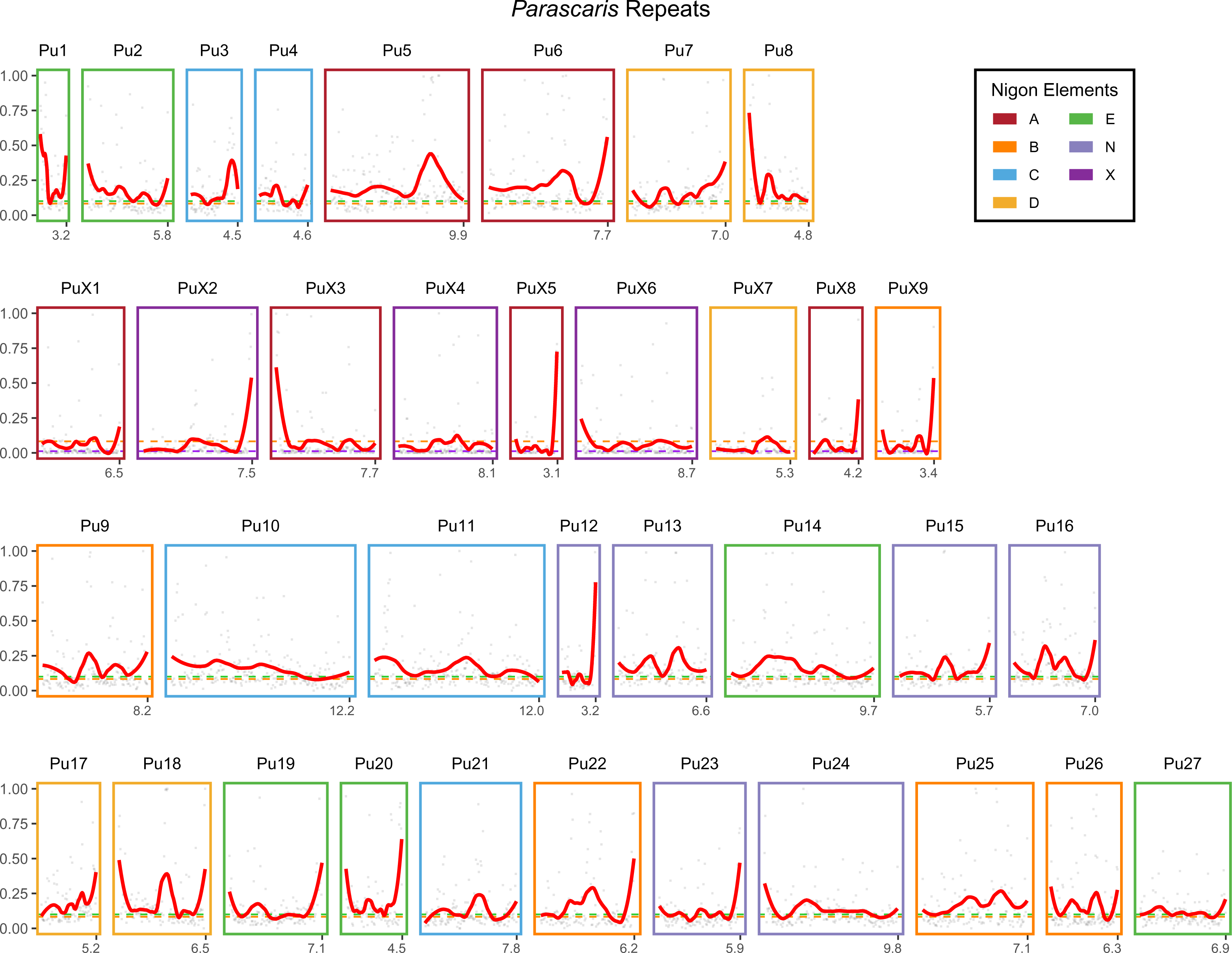
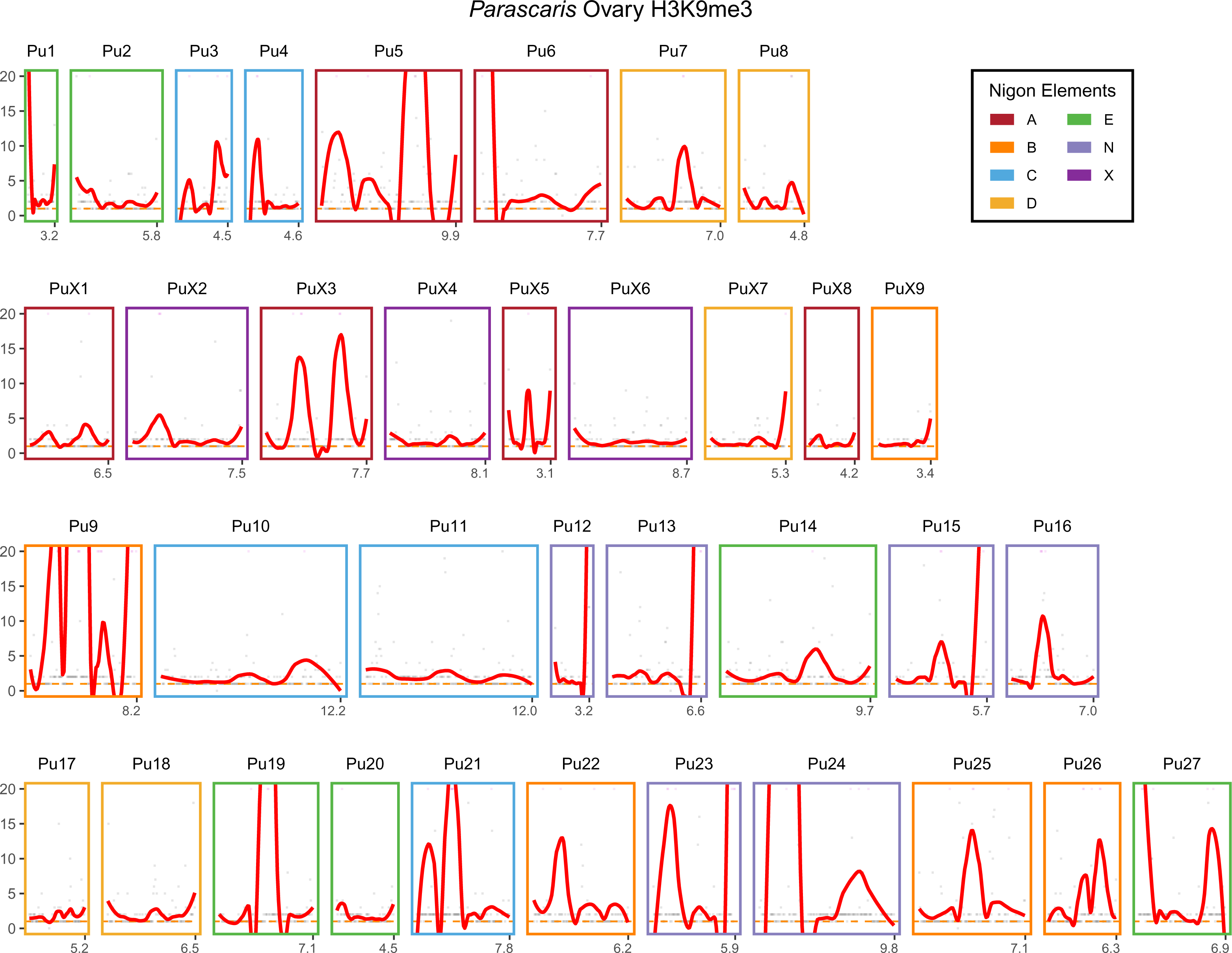
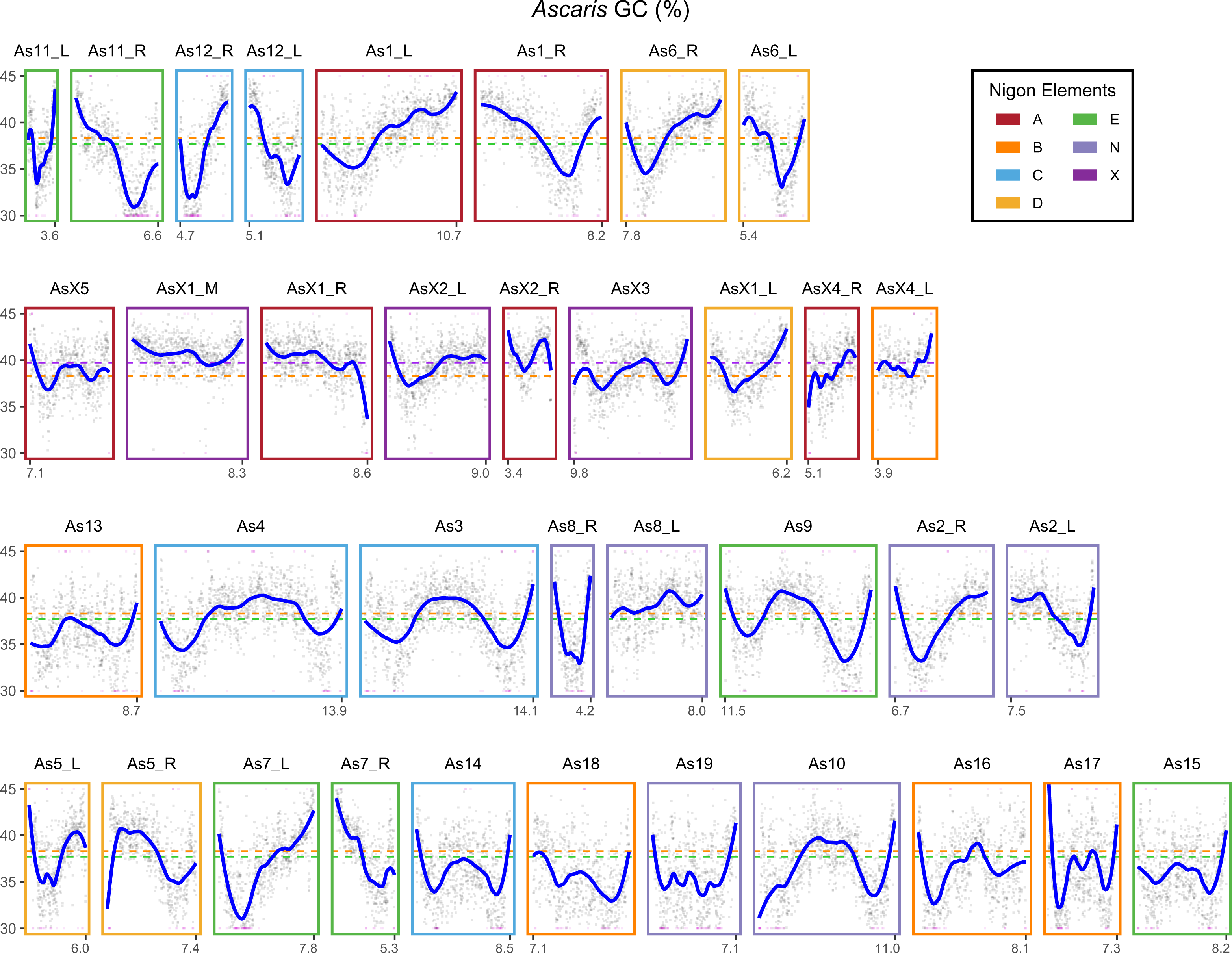
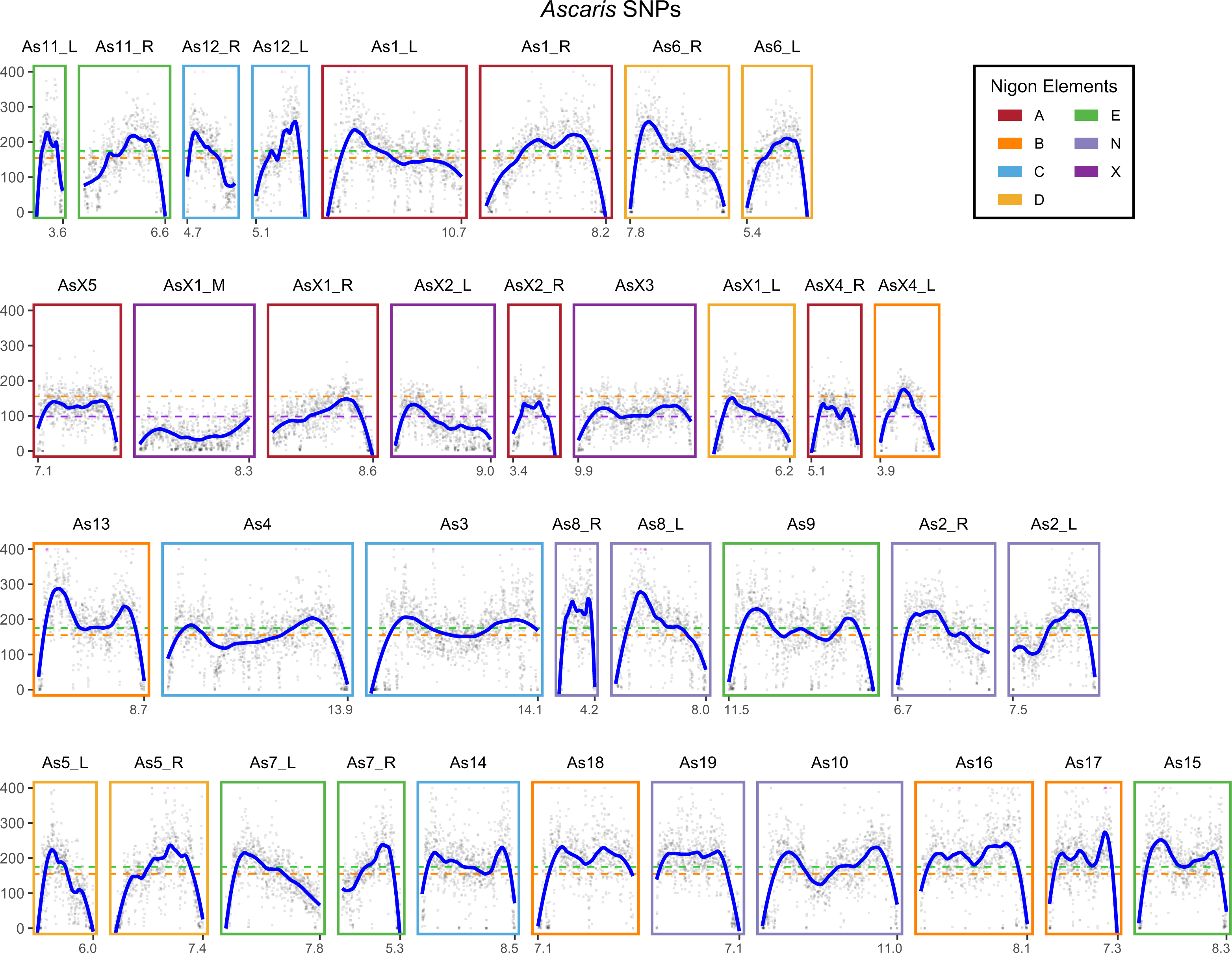
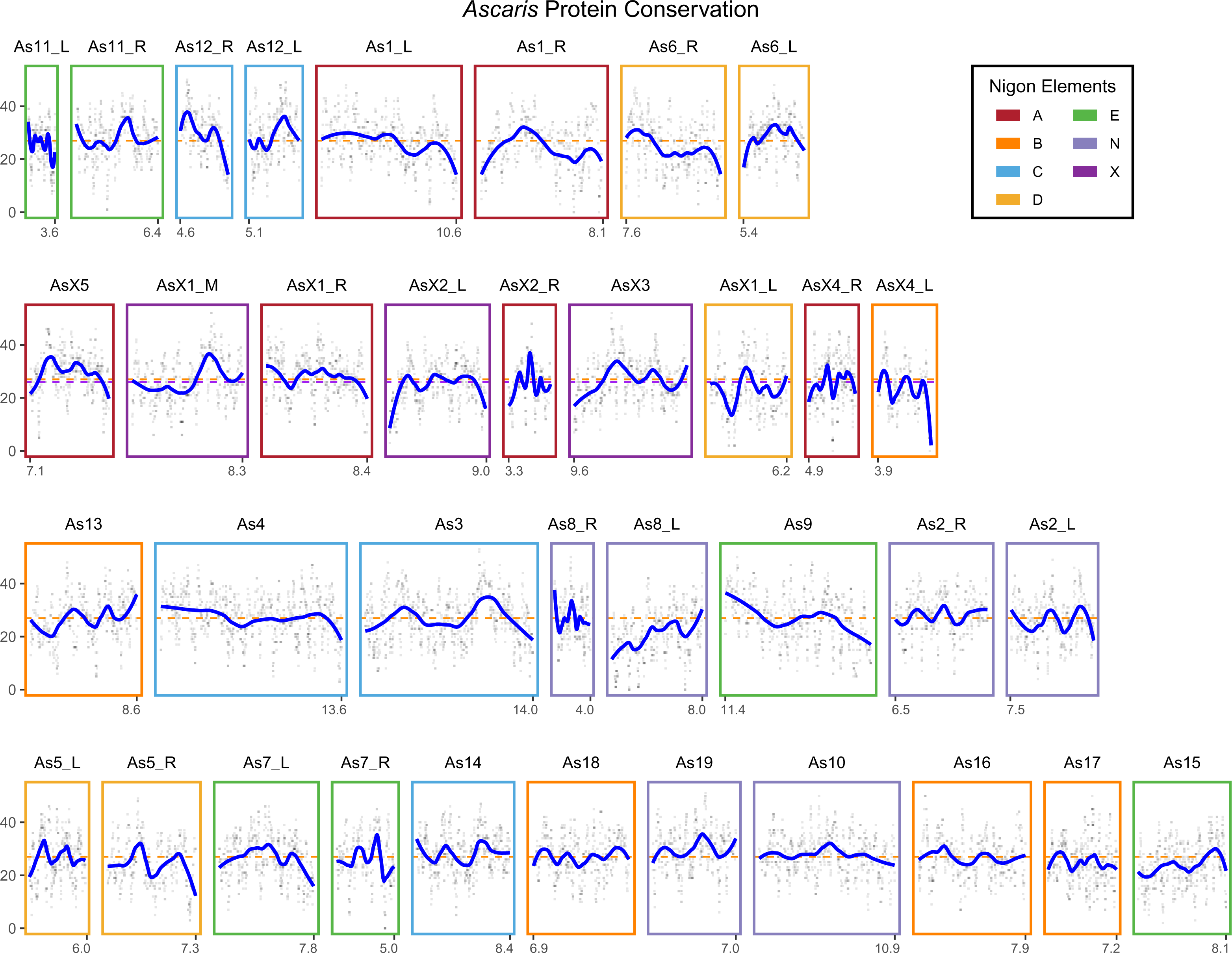
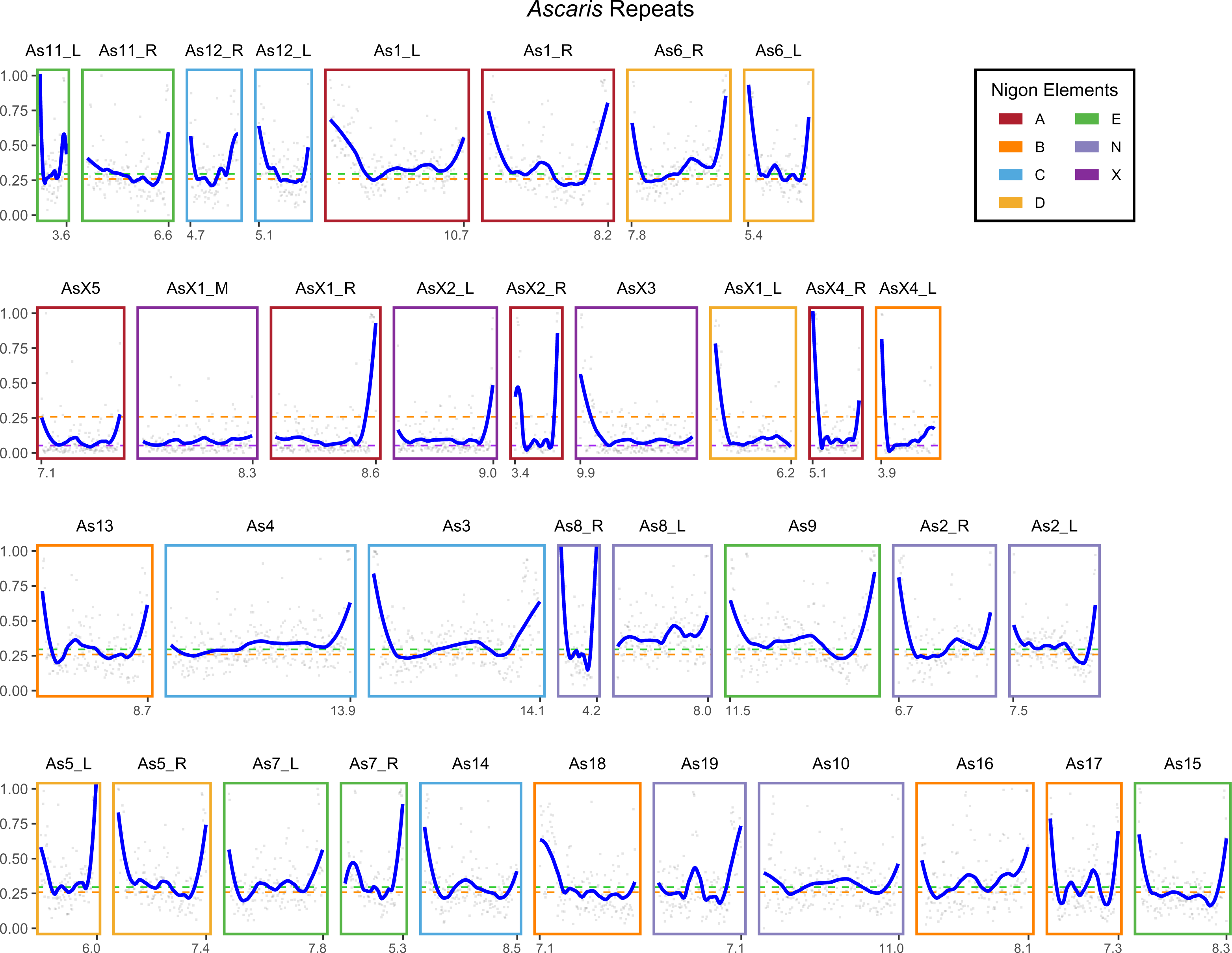
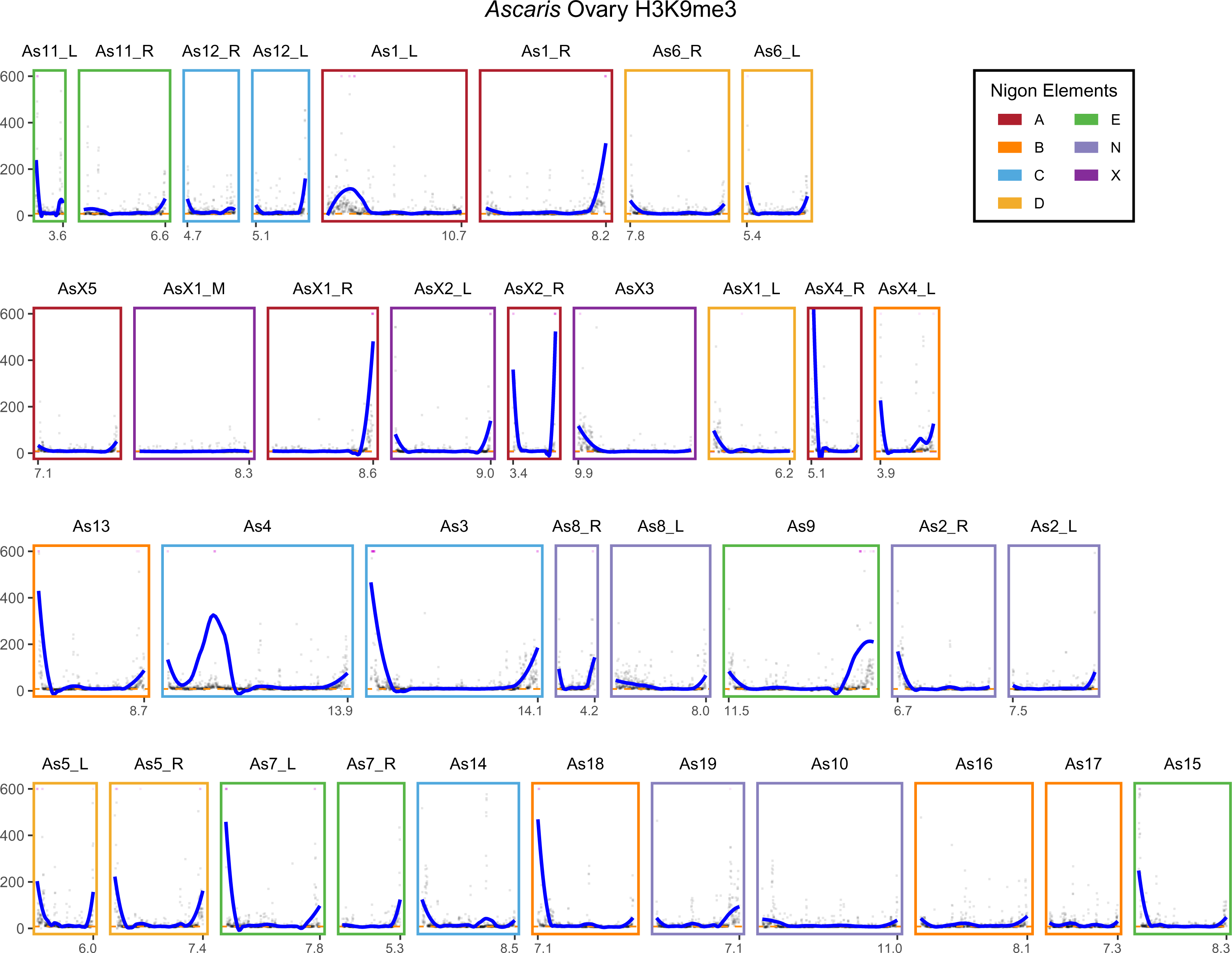
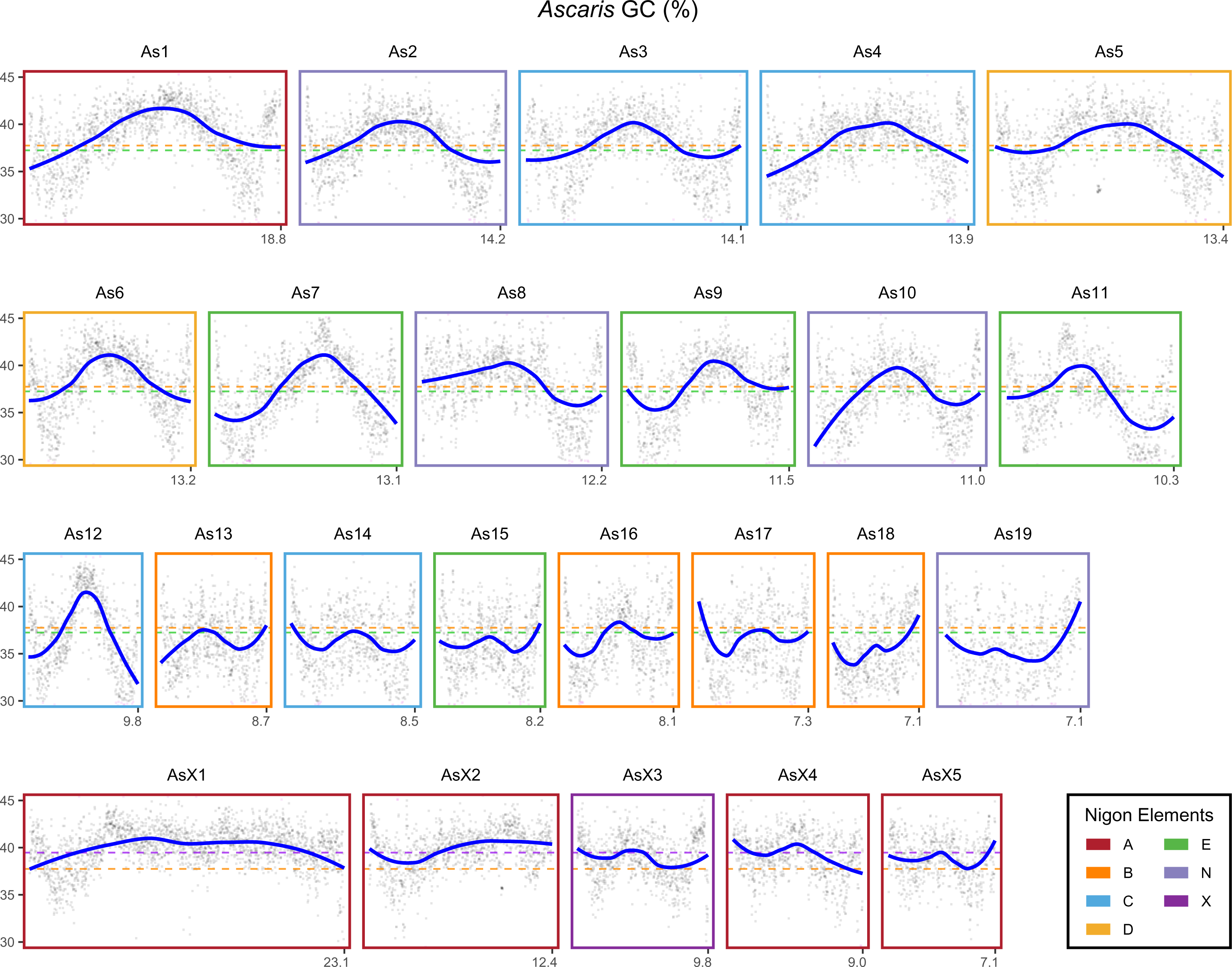
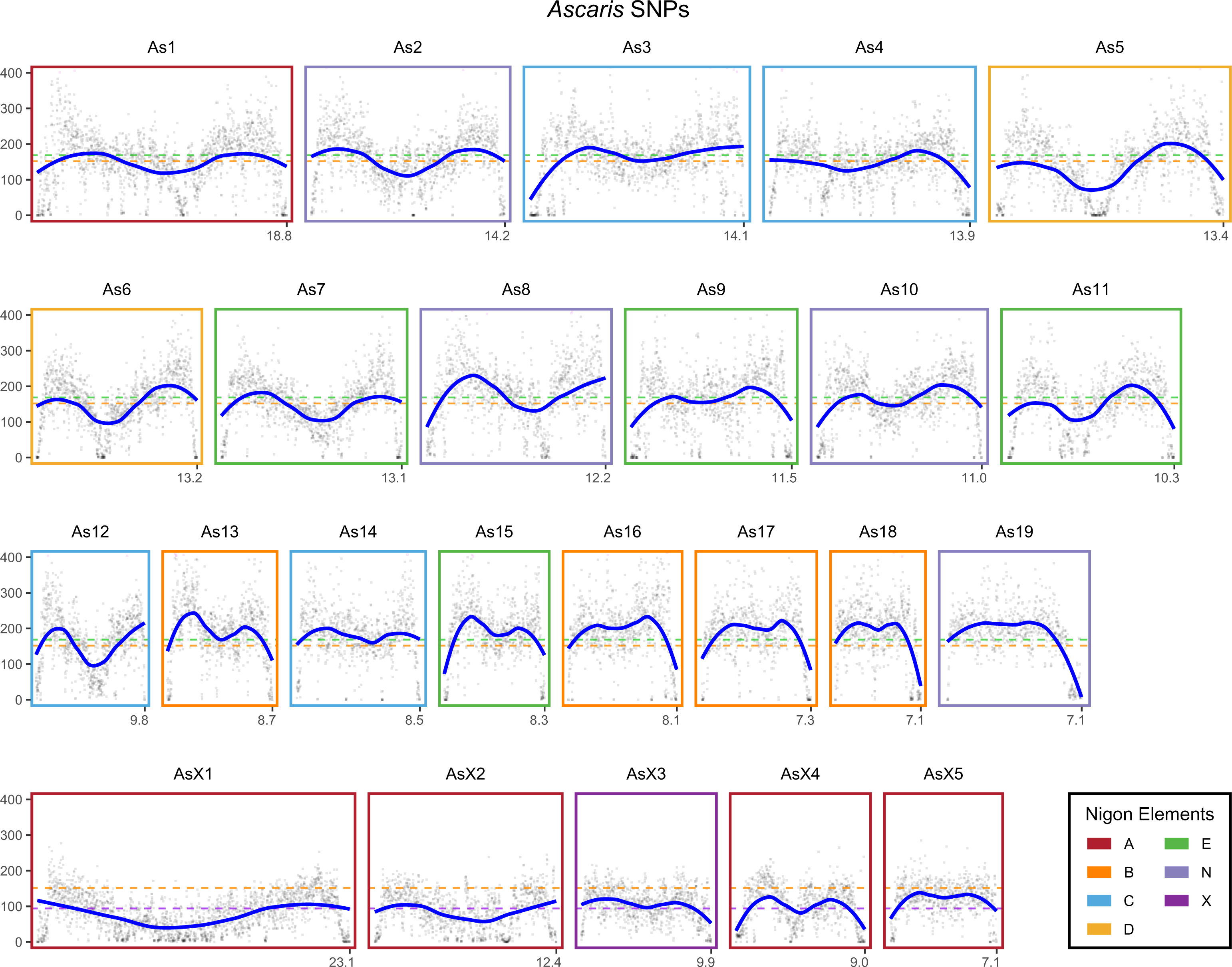
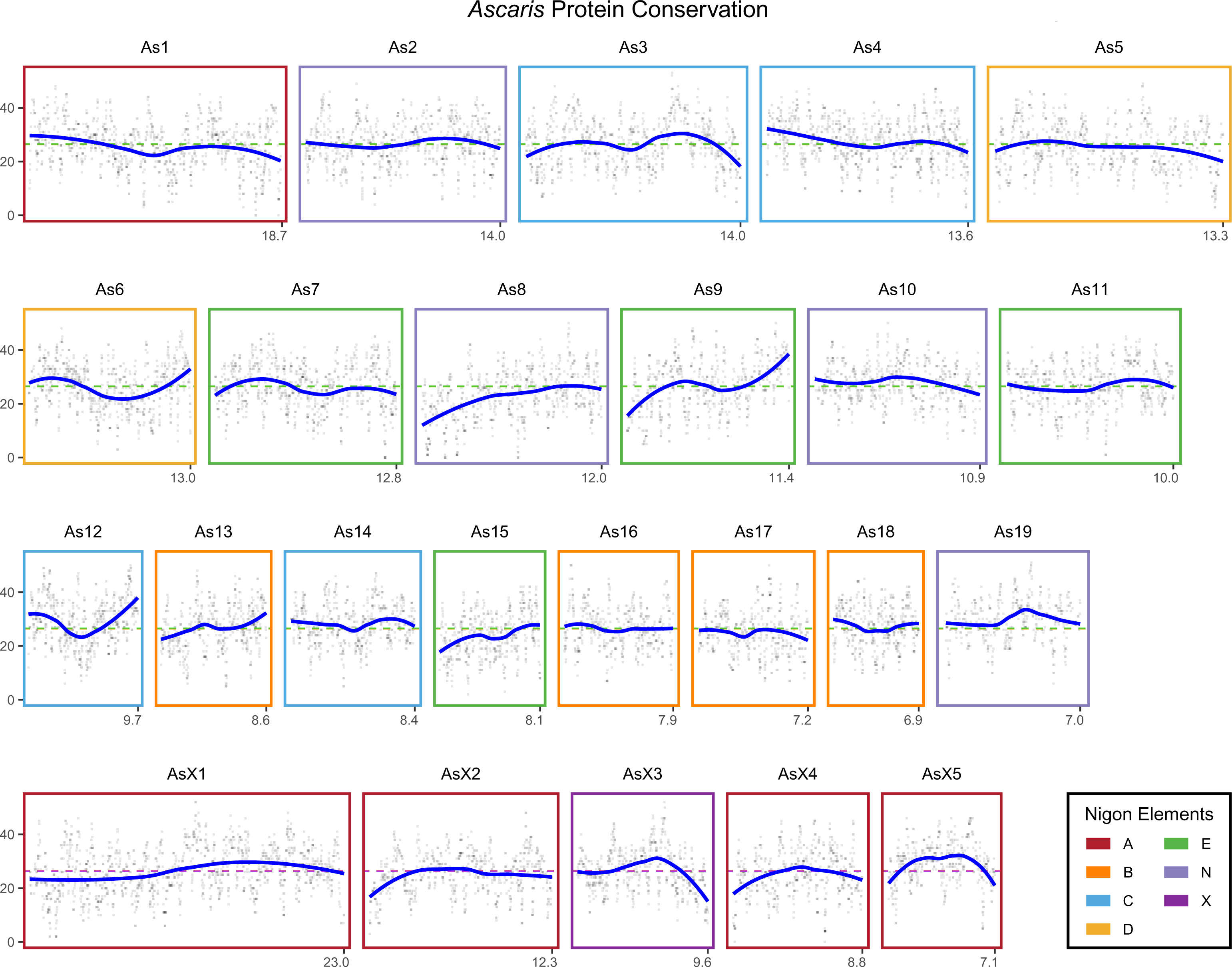
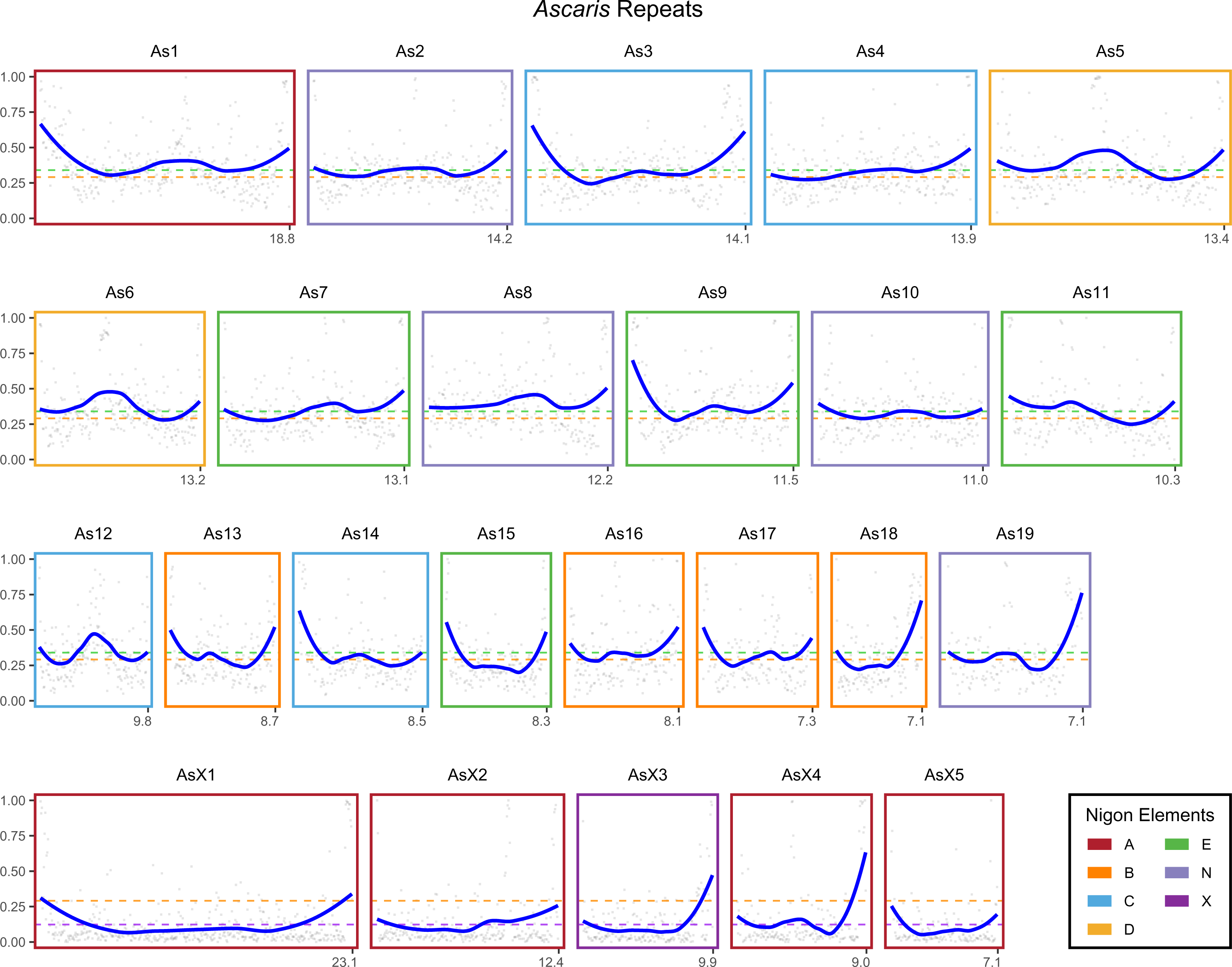
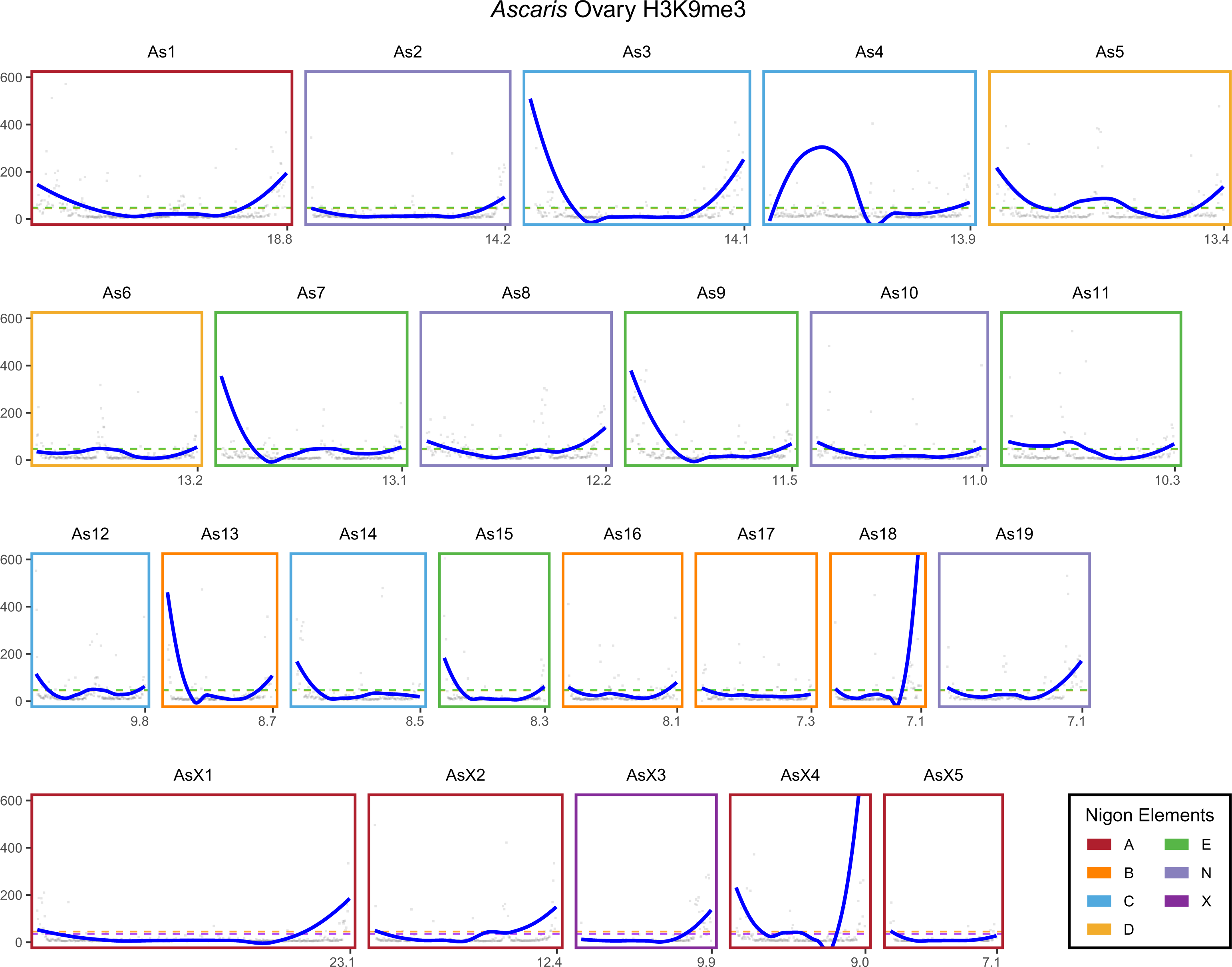
Genomic features for chromosomes in *Parascaris* and *Ascaris*. Features of GC content (%), SNP density, protein conservation, repeat density, and H3K9me3 (from the ovary) were plotted in 36 presumptive ancestral pre-fused chromosomes from *Parascaris* (the first five pages) and *Ascaris* (the next five pages) and the 24 germline chromosomes of *Ascaris* (the last five pages).

**Figure S4.**
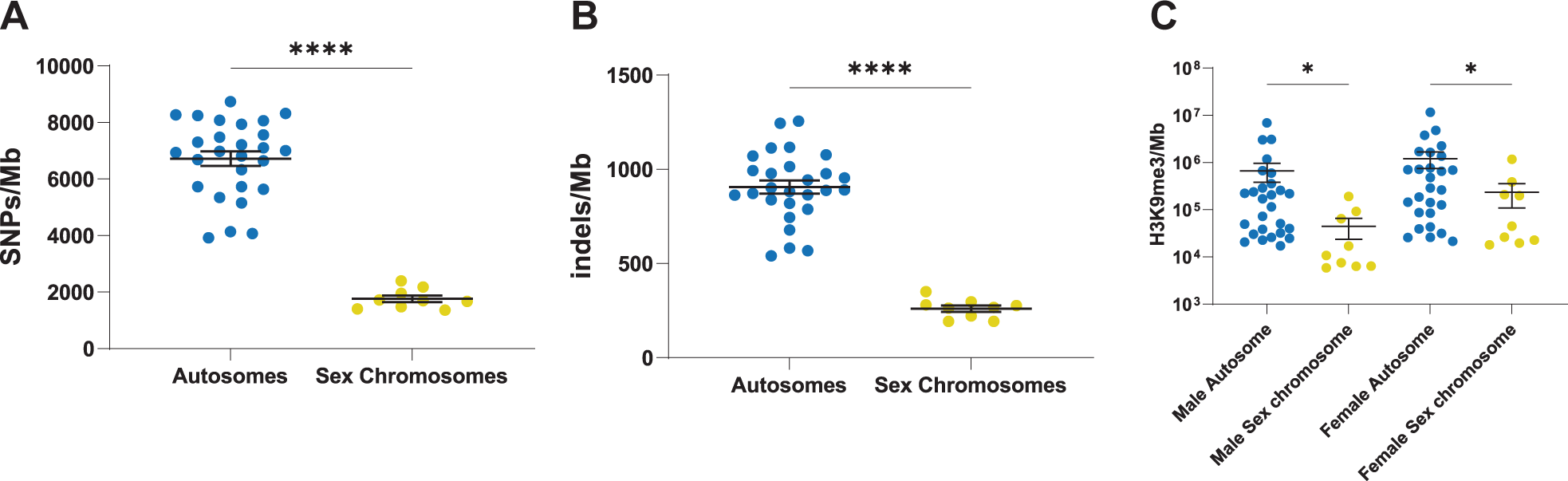
The region for *Parascaris* sex chromosomes lacks mutations and heterochromatin. Differences in SNPs, indels, and H3K9me3 are shown between the genomics regions that correspond to *Parascaris* somatic autosomes and sex chromosomes. P-values were determined using Welch’s t-tests (* P < 0.05, **** P < 0.001).

### Supplemental tables

**Table S1. *Parascaris* genes, annotation, and expression**

**Table S2. Synteny among ascarid genomes**

**Table S3. Nigon elements in Parascaris, Ascaris, and Baylisascaris**

**Table S4. Comparison of meiosis-related genes in nematodes**

### Supplemental movie

**Movie S1: 3D images of *Parascaris* germline chromosomes during spermatogenesis**

## Acknowledgments

We thank Martin Nielsen for the *Parascaris* material; graduate student Ruwaa Mohamed for initial PacBio data analysis; undergraduate students Eduardo Villalobos and Abigail West for helping with staining; and Rachel Patton McCord for interpretation of the Hi-C data. We also thank Tom Dockendorff and other lab members for helpful discussions and Mariano Labrador, Albrecht von Arnim, and Dick Davis for comments and critical reading of the manuscript. This work was supported by NIH grants to J. W. (AI155588 and GM151551) and the University of Tennessee Knoxville Startup Funds.

## Author contributions

J.R.S. and J.W. designed the project; M.V.Z prepared the genomic DNA for PacBio and did RNA-seq; J.R.S. performed Hi-C and staining; R.O. and J.R.S. did CUT&RUN; J.R.S. did imaging analysis; J.R.S., B.E., S.B.Z., M.V.Z., R.O., and J.W. carried out bioinformatic analyses, with J.R.S. and J.W. assembled the genome, S.B.Z. defined gene models, M.V.Z. identified repeats, B.E., J.R.S., and J.W. analyzed the genomic features and carried out comparative genomics analysis, B.E., R.O., and J.W. analyzed CUT&RUN data; and J.W. wrote the manuscript with assistance from J.R.S. and B.E.

